# Abundance and distribution of ice-associated seals in the western Bering Sea and Sea of Okhotsk, 2012-2013

**DOI:** 10.64898/2026.09.03.749204

**Authors:** Paul B. Conn, Irina S. Trukhanova, Peter L. Boveng, Vladimir Chernook, Alexander N. Vasiliev

## Abstract

In the springs of 2012-2013, Russian Federation scientists conducted aerial surveys of ice-associated seals (including bearded, ribbon, ringed, and spotted seals) in the Sea of Okhotsk and in Russian waters of the western Bering Sea. These instrument-based surveys used a combination of thermal imaging and digital photography to detect and photograph seals basking on sea ice. Initial estimates of seal distribution and abundance from these surveys have been published in Russian journals, but did not make adjustments for observation error. In this paper, we reanalyze these data with the goal of producing spatially explicit estimates of seal abundance that properly account for incomplete detection and species classification errors. Our analysis used estimated relationships between seal abundance and habitat variables to predict abundance in unsurveyed locations, and auxiliary data on detectability to model the observation process. Under this framework, we estimated that in 2012 there were approximately 185,330 bearded seals, 68,564 ribbon seals, 272,609 ringed seals, and 226,003 spotted seals in the western Bering Sea; in 2013, these numbers were 144,105, 39,435, 203,130, and 90,501, respectively. In the Sea of Okhotsk, estimated abundance was 218,740 bearded seals, 221,228 ribbon seals, 179,893 ringed seals, and 169,870 spotted seals (2013 only). These are the first credible estimates of ice-associated seals in the Okhotsk and western Bering Seas, and should be useful for population management and for measuring how density and distribution change as a function of decreasing Arctic sea ice. Declines in seal estimates in the western Bering Sea from 2012 to 2013 mirror patterns observed in the eastern Bering Sea during concurrent U.S. surveys and could represent a decrease in abundance or a substantial decrease in pupping and molting activity.

## 1 Introduction

Despite broad concern for the viability of ice-associated marine mammals in the face of Arctic warming, little is known about current population trends for many species, including bearded (*Erignathus barbatus*), ringed (*Pusa hispida*), ribbon (*Histriophoca fasciata*), and spotted (*Phoca largha*) seals [Laidre et al., 2015]. These species are largely dependent on sea ice in boreal spring (April - June) for life history functions such as molting and pupping. Globally, average melt onset in the Arctic has been occurring earlier in the year than in decades past [Stroeve et al., 2014], with less ice present in the spring [Onarheim et al., 2018]. In the Bering and Okhotsk Seas, such trends have been less pronounced or non-existent up through 2017 [Stroeve et al., 2014, Onarheim et al., 2018], though climate projections suggest the quantity of spring ice, and thus crucial habitat for these species, will decrease in the future [Boveng et al., 2009, Cameron et al., 2010, Kelly et al., 2010, Boveng et al., 2013]. Indeed, 2018 and 2019 had extremely low spring ice extents in the Bering Sea, far surpassing previous records [Stabeno and Bell, 2019].

Monitoring trends of seal populations requires time series of absolute abundance or time series of relative abundance with consistent survey methodology. Although previous surveys have been conducted in the western Bering and Okhotsk Seas [Fedoseev, 2000], they have been either limited in spatial scope, poorly documented, or subject to biases of unknown magnitude. Further, platforms and methods used to survey seals have varied considerably so that it is difficult to compare estimates without first correcting for incomplete detection (i.e., estimating absolute abundance or density).

In the springs of 2012 and 2013, Russian and U.S. research teams conducted the first-ever comprehensive surveys of ice-associated seals (specifically, bearded, ribbon, ringed, and spotted seals) in the Bering Sea and Sea of Okhotsk. Each survey team used fixed-wing aircraft to count seals basking on sea ice, using thermal (infrared) cameras to detect the warm bodies of seals on the cold sea ice, and coordinated digital photography to obtain color images to confirm species identity. Since it was infeasible to cross into foreign airspace, the Russian team surveyed the western Bering Sea (wBS) and the Sea of Okhotsk (SO), while the U.S. team surveyed the eastern Bering Sea (eBS).

Initial estimates of seal distribution and abundance from the wBS and SO were recently published in Russian journals [Chernook et al., 2014, 2018]. In particular, the authors used species distribution models fitted to count data to produce spatially explicit abundance estimates of basking seals. However, there are several challenges to interpreting these estimates as published. First, they do not account for seals that were in the water and therefore unavailable to be detected by infrared sensors. Second, observations of seals were grouped into several categories, with pups recorded separately. As such, species-specific estimates are for age 1+ animals (a species-unspecific estimate of “pups” is also available). Third, since a large proportion of the seals detected by thermal cameras were not photographed by color cameras, estimates were based on apportioned counts, where unknown seal species were added to species-specific counts based on the observed species proportions; uncertainty in the count apportioning process was not propagated when calculating precision of abundance estimates. Fourth, there is considerable potential for species misidentification when examining photographs [McClintock et al., 2015], a complication that was not addressed in initial modeling efforts. All of these factors combined can be expected to result in biased, overly precise estimates of seal abundance.

Recently, the U.S. survey team produced estimates of abundance for ice-associated seals in the eBS in 2012 and 2013 [Boveng et al., 2026] and for subsequent surveys conducted in the Chukchi Sea in 2016 [Boveng et al., 2025]. They used models that accounted for changing spatial distributions over the course of the surveys, as well as many of the nuisance observation processes that affect seal counts (including incomplete detection and species misidentification; Conn et al. 2014). Although any modeling approach is imperfect, it is our view that abundances estimated in this manner are more robust for inference because many sources of bias are simultaneously accounted for. However, the methods of Boveng et al. (2025, 2026) cannot be directly applied to Russian surveys in the wBS and SO because of differences in how data were collected and summarized.

In this paper, we reanalyze Russian survey data in 2012 and 2013 with the goal of providing unbiased estimates of ice-associated seals in the wBS and SO. In particular, we analyze count and effort data provided in Russian journals [Chernook et al., 2014, 2018], incorporating auxiliary data from satellite tagging studies and detection experiments to better account for observation error and species misidentification. Importantly, we propagate uncertainty from these auxiliary data sources into abundance estimates. We also properly account for thermal detections of seals that don’t have accompanying color photographs.

## Methods

Before describing methods for modeling seal abundance from survey counts, we first provide a brief description of the study areas and aerial survey protocols. See Chernook et al. (2014,2018) for a more detailed description.

### Survey methods & study areas

In April and May of 2012 and 2013, a Russian survey team conducted aerial surveys over sea ice in the wBS and SO. Flights were intended to include all ice covered areas.The study area in the wBS consisted of all areas south of the Bering Strait and west of the U.S. Exclusive Economic Zone (EEZ) boundary that were covered by ice at some time during the springs of 2012-2013 (Fig. 1). Because spatial coverage was low in the SO in 2012, we only used flights flown in 2013 for abundance estimation in this area. The SO was defined as occurring within waters bounded by the Russian mainland to the north and east, by Hokkaido and the Kuril islands to the south, and by Sakhalin island to the west (Fig. 2).

**Figure 1:**
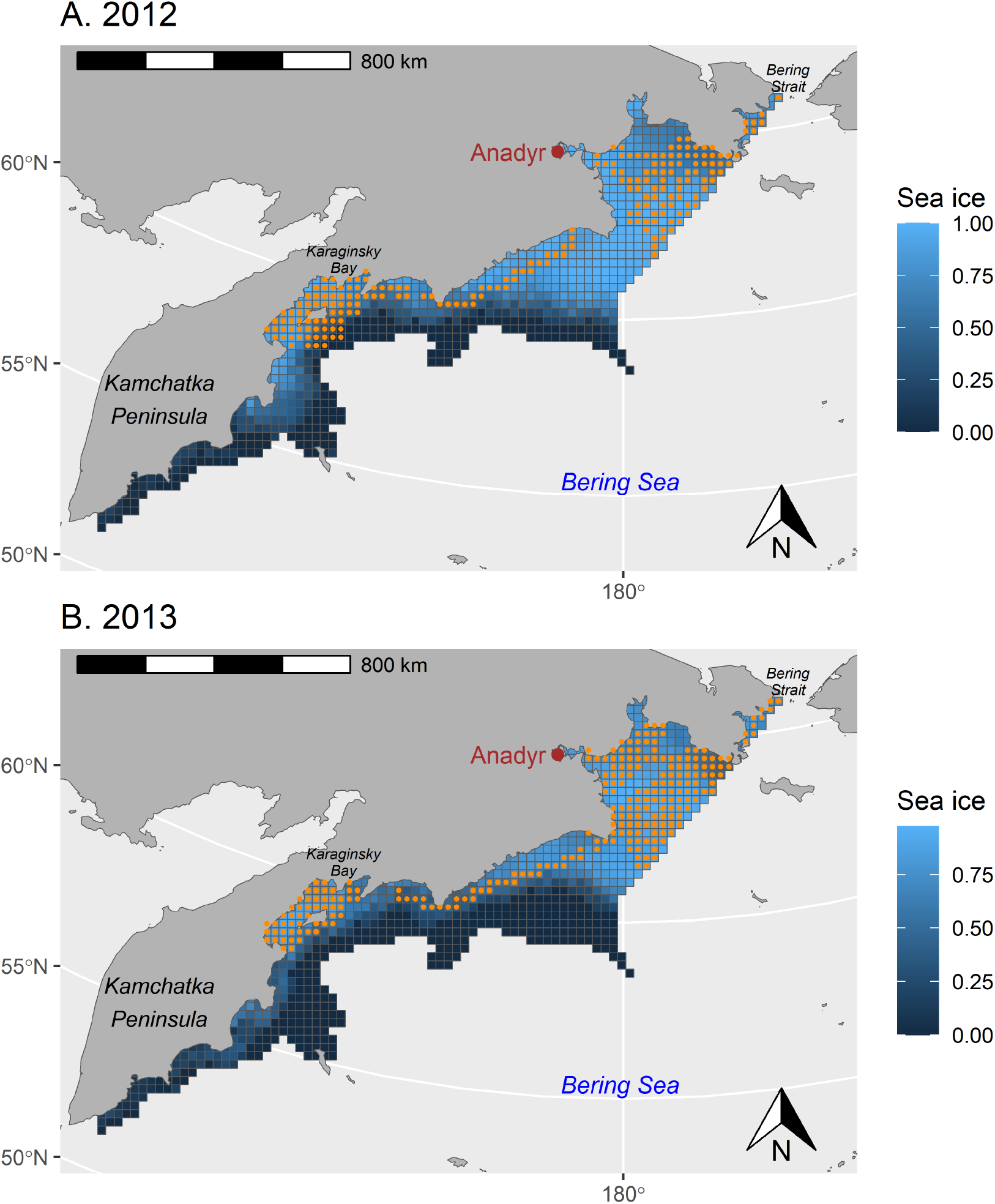
Study area and analysis grid for 2012 and 2013 aerial surveys of the western Bering Sea. Grid cells receiving survey effort are indicated in orange. Blue to black shading represents averaged remotely sensed proportional sea ice

**Figure 2:**
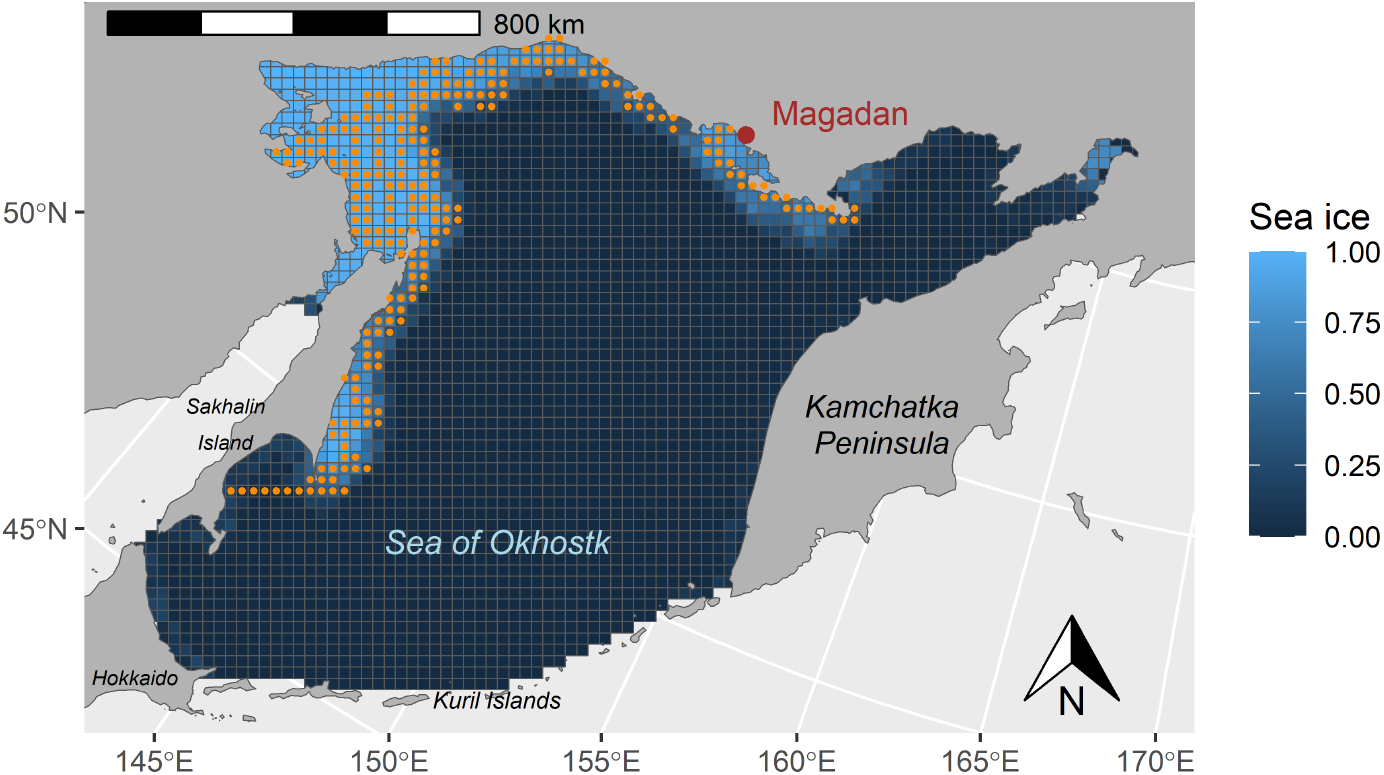
Study area and analysis grid for 2013 aerial surveys of the Sea of Okhotsk. Grid cells receiving survey effort are indicated with orange points. Blue to black shading represents remotely sensed proportional sea ice concentration. Each cell is *≈* 625 km^2^.

Flights were conducted with a fixed-wing aircraft flying at 200-250 m altitude and equipped with an infrared Malakhite-M scanner installed underneath the aircraft to detect the warm bodies of seals against the cold sea ice substrate with 0.2*^◦^*C sensitivity. Seals were simultaneously photographed with 3 Nikon D800 36 MP digital cameras (focal distance 50 mm) to determine species identity whenever possible. The effective area sampled was calculated from altitude, focal length, and camera angles to determine the footprint of the thermal scanner, and then using standard distance sampling models [Buckland et al., 2001] to calculate an effective strip width [Chernook et al., 2014, 2018]. The latter procedure was used because lower densities of seals were detected at the periphery of the thermal footprint than in the center, suggesting that detectability of the thermal camera decreased slightly with distance from the transect line.

In 2012, the Russian survey team flew 7224 km of transects in the wBS, effectively covering 3374 km^2^. In 2013, they flew 6737 km of transects in the wBS, effectively covering 3881 km^2^ (Fig. 1). Also in 2013, 5617 km of transects were flown in the SO, effectively sampling a 2993 km^2^ swath (Fig. 2). Okhotsk flights were all conducted within a relatively short time window (1 May - 9 May) in 2013. Flights in the wBS were conducted over a longer period: 20 April - 5 May in 2012 and 11 April to 30 April in 2013.

To estimate abundance, we used effort and observations summarized within 25*×*25 km^2^ grid cells reported in Russian journals; exact plots of transects flown are available in the original articles [Chernook et al., 2014, 2018]. Within each grid cell, data were summarized as follows: area surveyed, total numbers of seals detected via infrared, number of age 1+ seals photographed by species (ringed, bearded, spotted, ribbon, or unknown) and number of pups photographed.

### Predictive covariates

We obtained covariate data to help explain variation in survey counts and to predict abundance in unsampled locations. We downloaded remotely sensed sea ice concentration data from the National Snow & Ice Data Center, Boulder, Colorado, USA at an approximate 25 *×* 25 km grid cell resolution for the date ranges in our surveys. We used a daily resolution for the wBS, but averaged values over the short (9-day) duration of the survey for the SO. We used these covariates directly in predictive models, and also to calculate distance from each grid cell centroid to the ice edge (for each day of the survey in case of the wBS). We also obtained bathymetry data to calculate the water depth at the centroid of each grid cell using the online Grid Exact tool produced by NOAA National Centers for Environmental Information (https://maps.ngdc.noaa.gov/viewers/wcs-client/) and to calculate the distance to the continental shelf break. For the wBS, we defined the shelf break to be the 1000 m isobath, a feature known to be important for upwelling and primary productivity. The SO is much shallower and we defined “shelf break” to occur at the 200m isobath. Land mass data were obtained from the ptolemy package [London, 2018] to calculate distance to land. For the wBS, we followed the approach of Chernook et al. (2018), modeling the effects of covariates differently for three spatial strata (Northern: *>* 62.4*^◦^N*; Southern: *<* 167.8*^◦^E*; Middle: Otherwise). All calculations were made in the R programming environment [R Development Core Team, 2017] with frequent use of the sp and rgeos packages [see e.g. Bivand et al., 2013]. Absolute correlations between continuous predictive covariates were all *<* 0.55.

### Auxiliary detection data

A variety of processes conspire to prevent straightforward interpretation of seal detections obtained in aerial surveys, including:

1. *Availability bias*– not all individuals are “available” to be detected on sea ice since some are in the water;
2. *Escape behavior* – Some individuals escape into the water prior to being detected;
3. *Perception bias*– Thermal sensors may miss some individuals even if they are on ice;
4. *Species classification*– Not all thermally detected individuals are photographed, no species information is available for pups, and observers sometimes misclassify seals when making species determinations from photographs.

We next summarize auxiliary data available for addressing each of these potential sources of bias.

### Availability

To account for unavailable animals, we used data from seal-borne satellite-linked wet/dry bio-loggers to predict the proportion of animals that are basking on ice in conditions similar to those observed in the field. Bio-logger records for bearded, ribbon, and spotted seals deployed between 2005 and 2020 in the Bering and Chukchi Seas were previously analyzed by London et al. (2024), who demonstrated the importance of effects such as age-sex class, day-of-year, time-of-day, and weather covariates when predicting haul-out behavior. For the wBS, we used the models of London et al. (2024) to predict availability probability for species *k* with space-time index *j*, *a_k,j_* for these three species. We summarized uncertainty in predictions with a variance-covariance matrix Σ*_k_* on the logit scale. To make predictions, we first downloaded weather data from the North American Regional Reanalysis [NARR; Mesinger et al., 2006] for the locations and times when surveys were conducted. The NARR data were unavailable for the SO, so we instead made predictions using haul-out models that omitted weather effects (obtained by rerunning the models of London et al. (2024) and removing weather covariates).

Until recently, availability correction factors used in aerial surveys for ice-associated Arctic seals [e.g., Bengtson et al., 2005, Conn et al., 2014, Ver Hoef et al., 2014] ignored any age- and sex-based variation in haul-out probabilities. The implicit assumption was that the behavior of telemetered animals represents the population as a whole. However, if haul-out probabilities are related to sex- and age-class, and if there are tangible differences between the age- and sex-structure of the telemetered sample and that of the population, correction factors computed in this way may lead to biased abundance estimates [Conn and Trukhanova, 2023]. This finding recently led Boveng et al. (2025) to calculate availability as a weighted average of predictions from each age- and sex-class. Although the sample size of bearded seals with bio-loggers was too small to permit reliable estimation of age- and sex-effects on haul-out probabilities, and ringed seal availability predictions lack age- and sex-specificity [Boveng et al., 2026], we used age- and sex-weighting for ribbon and spotted seals. In particular, we computed

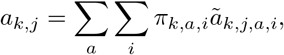

where *ã_k,j,a,i_* is the predicted haul-out probability for age class *a* (pup, subadult, or adult) and sex *i* seals of species *k* at space-time index *j*. The *π_k,a,i_*values represent population-level proportions of seals of species *k* that are of age class *a* and sex *i*. We set *π_k,a,i_* equal to stable stage proportions calculated from survival, fecundity, and maturity schedules [Conn and Trukhanova, 2023] and used the delta method [Dorfman, 1938] to calculate the associated variance-covariance matrix for weighted predictions. Predictions of availability for bearded seals, ribbon seals, and spotted seals were generated using the highest ranked models from London et al. (2024), with environmental covariates particular to our study.

So far, we have described how we modeled availability of bearded, ribbon, and spotted seals, for which haul-out probabilities are equivalent availability. Ringed seals are potentially more problematic, especially in regions where they can be hauled out but still concealed within snow dens [subnivean lairs; McLaren, 1958, Smith and Stirling, 1975]. In the SO, ringed seals are not believed to use subnivean lairs, or at least if they do it is at substantially reduced frequency [Fedoseev, 1965, 1975]. However, ringed seals in the wBS likely include seals that construct lairs in landfast ice as well as those that haul out and have pups on pack ice without snow lairs [Fedoseev, 2000]. Recent analyses based on temporal trends in aerial survey counts in the Chukchi Sea suggested that the proportion of visible seals increases as snow melts (which leads to lair collapse) [Lindsay et al., 2021, Boveng et al., 2025], but this relationship is less clear in the Bering Sea [Lindsay et al., 2021, Boveng et al., 2026]. Our preference in this paper is to follow Boveng et al. (2026) who conservatively assumed that all hauled out ringed seals are visible to aircraft while surveys are conducted. This approach will likely lead to some negative bias, but resulting “minimum” estimates may remain useful for management [Boveng et al., 2026]. We investigate sensitivity of abundance estimates to alternative forms of ringed seal availability in *Sensitivity analyses*.

### Escape behavior

Estimates of availability probability measure the expected proportion of seals that are potentially visible to aircraft in natural (i.e., undisturbed) conditions. However, if some seals exhibit behavioral disturbance to aircraft, there will be an additional fraction that go undetected because they escape into water before the thermal scanner passes overhead. To quantify disturbance, we used data on disturbance experiments reported in other studies. We based bearded and ringed seal disturbance rate on disturbance trials conducted by Russian survey personnel on similar flights over the Chukchi Sea [Boveng et al., 2025], and ribbon and spotted seal disturbance rates based on U.S. trials conducted during eBS surveys [Boveng et al., 2026]. In each case, forward facing trials were used to estimate the proportion of each species that escape into water before entering the range of thermal cameras. Escape proportions and standard errors (SE) for bearded, ribbon, ringed, and spotted seals were 0.06 (0.03), 0.0 (0.0), 0.30 (0.03), and 0.04 (0.04), respectively. We use these estimates and associated standard errors in current modeling efforts as they were the best data available (no such exercise was conducted in the SO or wBS surveys). However, we note that the aircraft used by the Russian survey team was larger, louder, and flew at a lower target altitude than the aircraft used in U.S. surveys, factors which might be expected to increase disturbance rates for ribbon and spotted seals relative to the values here. Conversely, the Russian aircraft was much faster than that used in U.S. eBS surveys, so there was less time for seals to respond before entering the radius of the thermal detector.

### Perception bias

Previous analysis of count data from 2012-2013 Russian seal surveys [Chernook et al., 2014, 2018] accounted for incomplete detection of thermal sensing equipment via effective strip width calculations typical of distance sampling applications [Buckland et al., 2001]. This approach implicitly assumes that infrared sensors detect 100% of the seals on ice directly under the aircraft (i.e., at distance zero from the transect line), an assumption that seems reasonable for these surveys. For instance, using a double sampling experiment, Boveng et al. (2026) reported thermal detection rates of 94-97% in U.S. survey data. Russian surveys were conducted with a more powerful sensor at a lower altitude, suggesting that detection probability directly beneath their aircraft was close to 100% in good survey conditions. We make this assumption in subsequent modeling efforts.

### Species classification

Trukhanova et al. (2018) analyzed data from a double observer experiment applied to images taken in Russian surveys using methods described by McClintock et al. (2015). Their analysis revealed moderate to high levels of species misidentification; marginalizing over certainty category (an additional variable needed for running such an analysis) and limiting results to the primary observer yielded misclassification rates as high as 23% (Table 1). We use these estimates, together with an associated variance-covariance matrix in multinomial-logit space, to model species misidentification and propagate uncertainty in subsequent modeling efforts.

**Table 1:** Estimates of species classification probabilities used in analysis of seal survey data. Given a true species, entries give probabilities of obtaining one of five types of species classification (including ‘unknown species’). Estimates are for the primary observer, and were calculated directly from output produced by Trukhanova et al. (2018).

| True species | Observation type |  |  |  |  |
| --- | --- | --- | --- | --- | --- |
|  | Spotted | Ribbon | Bearded | Ringed | Unknown |
| Spotted | 0.78 | 0.01 | 0.01 | 0.13 | 0.08 |
| Ribbon | 0.00 | 0.97 | 0.00 | 0.01 | 0.01 |
| Bearded | 0.05 | 0.06 | 0.77 | 0.06 | 0.06 |
| Ringed | 0.01 | 0.01 | 0.01 | 0.96 | 0.01 |

### Statistical models for seal counts

For both study areas (SO and wBS), our goal was to formulate models that related counts to (i) underlying levels of seal abundance and (ii) nuisance observation processes. In particular, we used a Tweedie error distribution [Jørgensen, 1987, Peel et al., 2013] to relate observed and expected counts, which allows for overdispersion relative to commonly used distributions for count data (such as the Poisson) and simultaneously accommodates zero inflation [excess zeroes relative to those predicted by standard probability distributions; Agarwal et al., 2002]. Let *C_j,o_* denote the observed count obtained in a grid cell with space-time index *j* (*j ∈* 1, 2*, · · ·, J*), and observation class *o* (*o ∈* 1, 2*, · · ·,* 7) corresponding to spotted seal, ribbon seal, bearded seal, ringed seal, photographed but unknown seal species, pup, and unphotographed seal. The final category is for cases where seals are thermally detected but there are no associated color images. Similarly, let *λ_j,o_* = *E*(*C_j,o_|**θ**,* **D**) denote an expected count given a vector of unknown parameters, ***θ***, and additional data on detection (e.g., area surveyed), **D**. Under this framework, inference proceeds by conditioning on observed counts and maximizing the likelihood

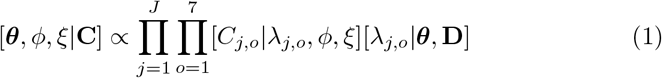

relative to *ϕ*, *ξ*, and ***θ***. Here, [*C_j,o_|λ_j,o_, ϕ, ξ*] specifies a Tweedie distribution with mean *λ_j,o_* and variance *ϕ_o_λ^ξo^*, evaluated at the observed data, *ϕ_o_* and *ξ_o_* are unknown variance parameters, and [*λ_j,o_|**θ***] is the conditional distribution of *λ_j,o_* given additional parameters ***θ*** (i.e., abundance, regression, and detection parameters). In subsequent applications, we restrict *ϕ_o_ >* 0 and 1 *< ξ_o_ <* 2, resulting in a version of the Tweedie known as a compound Poisson-gamma distribution [Lecomte et al., 2013] which has been shown to fit marine mammal data well in other applications [e.g., Sigourney et al., 2020, Boveng et al., 2025].

Our statistical models tabulate expected counts in each grid cell, *λ_j,o_*, as a function of unknown parameters representing abundance and detection. Using vector notation, and defining ***λ****_j_* = [*λ_j,_*_1_ *λ_j,_*_2_ *· · · λ_j,_*_7_]*^′^*, we write

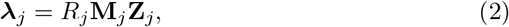

where *R_j_* is the proportion of sea ice habitat that is surveyed in the cell with spatio-temporal index *j*, **Z***_j_* = [*Z*_1*j*_ *Z*_2*j*_ *Z*_3*j*_ *Z*_4*j*_]*^′^*denotes latent abundance of spotted, bearded, ribbon, and ringed seals, respectively, and **M***_j_* is a (7 *×* 4) matrix operator encompassing confusion and thinning processes. We structure the elements of **M***_j_*, *m_j,o,k_* as follows:

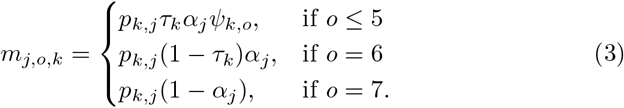

Here, *p_k,j_* defines a composite probability of detection for species *k* when surveying a grid cell with spatio-temporal index *j*, *τ_k_* gives the proportion of the population of species *k* that is composed of non-pups (here assumed time constant), *α_j_* is the proportion of seals that are photographed (assumed known and set to empirical values from the field), and *ψ_k,o_* gives the probability of classifying an adult of species *k* into observation type *o* given that it is photographed.

In practice, availability bias, escape behavior, and perception bias all contribute to detectability that is less than 1.0. We thus write detection probability as

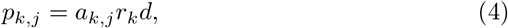

where *a_k,j_* gives the probability a seal of species *k* is available to be detected, *r_k_*is the probability an available seal remains on ice as the aircraft flies overhead, and *d* gives the probability of perception by the thermal sensors. However, as described previously, we set *d* = 1.0 for these surveys because distance effects were handled through effective strip width calculations and perception directly under the aircraft is thought to be close to 100%.

We use slightly different approaches to model variation in latent abundance, **Z***_j_*, depending on the analysis type. In particular, surveys in the Sea of Okhotsk were conducted over a relatively short, 9-day interval, and we found it simplest to average over sea ice conditions encountered during surveys (i.e. ignoring time). By contrast, wBS surveys occurred over longer intervals with substantial change in sea ice conditions while surveys were conducted. In this case, we specified a model that allowed seal abundance to change over space and time.

Until now, we have used the single subscript *j* to index sampled times and locations. In making inference to a larger grid (or time series of such grids), it is necessary to consider a fuller process model. For the SO, we specify a process model for latent abundance, *µ_k,s_*, for species *k* and grid cell *s ∈* (1, 2*, · · ·, S*). In this case the connection between *Z_k,j_* and *µ_k,s_*is simply **Z***_k_* = **W**_1_***µ****_k_*, where **W**_1_ is an (*S ×J*) matrix with entries *w_s,j_* = 1 if the *j*th sampled cell has spatial index *s* and zeroes elsewhere. In essence, **W**_1_ selects the sampled cells from the list of total cells. For the wBS, we use a spatio-temporal process model where we specify latent abundance by species (*k*), spatial cell (*s*) and survey day (*t*), *µ_k,s,t_*. In this case, define **W**_2_ to be an *ST × J* matrix where *w*_(*t−*1)*S*+*s,j*_ = 1 if the *j*th sampled cell was grid cell *s* at time *t* and zero otherwise. Then forming the matrix

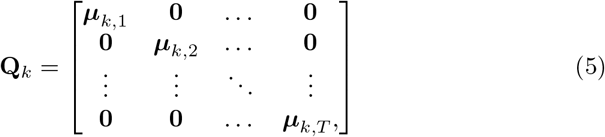

where **0** is a length *S* column vector of zeroes, and ***µ****_k,t_* = [*µ_k,_*_1*,t*_ *µ_k,_*_2*,t*_ *· · · µ_k,S,t_*]*^′^*, we have **Z***_k_* = **W**_2_**Q***_k_*. Again, we are simply picking the cells and times that are sampled out of a larger set.

For the SO, we modeled latent abundance for species *k*, ***µ****_k_* = [*µ_k,_*_1_ *µ_k,_*_2_ *· · · µ_k,S_*]*^′^*, as

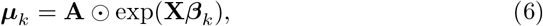

where **X** represents an (*S × b*) design matrix as typically used in generalized linear models [McCullagh and Nelder, 1989], ***β****_k_* represent a species-specific column vector of *b* regression parameters, **A** is a length *S* column vector giving the proportion of sea ice habitat in each grid cell, and ⊙ specifies product-wise multiplication (i.e., a Hadamard product). Under this framework, estimates of total abundance for the whole study area can be derived as *N*^^^*_k_* = Σ*_s_ µ_k,s_*.

For the wBS, we followed recommendations by Conn et al. (2015b) to stabilize estimation by including a single abundance parameter for each species, *N_k_*, that represents total abundance in the surveyed area, and assumed it was constant for the duration of each survey. We then let latent abundance values redistribute themselves on each day of the survey as ice conditions changed. Specifically, we modeled a length *S* vector of spatially-referenced latent abundances of species *k* on day *t*, (***µ****_k,t_*), as

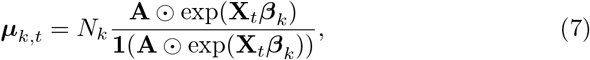

Here, the **1** represents a (length *S*) row vector of ones such that the denominator of Eq. 7 is a scalar sum of the numerator values (ensuring that Σ*_s_ µ_k,s,t_*= *N_k_*). Analagous to using a multinomial logit link function, there is no intercept in the design matrix **X***_t_* with this formulation.

### Data analysis & computing

We used different combinations of covariates for each data set to predict seal counts. We fit preliminary models with a relatively large number of predictors for each species, sequentially reducing model structure until we were able to satisfy numerical optimization convergence criterion (through application of the nlminb function in R; R Development Core Team [2017]) and achieve model predictions that reasonably fitted survey counts without obvious extrapolations in unsurveyed cells (e.g., predicted counts that were orders of magnitude greater than those observed during surveys). We used this strategy to identify a “general” model (sensu Burnham and Anderson [2002]) for each dataset. For the SO, models included sea ice concentration (linear and quadratic effects), distance from land, distance from shelf break, distance from ice edge^0.5^, and depth^1*/*3^ (linear and quadratic).

We specified Bayesian prior distributions on regression coefficients,

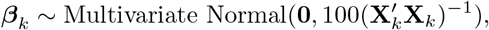

and on the logit of pup proportions:

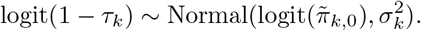

Here, *π^-^_k,_*_0_ gives mean pup proportions predicted from Conn and Trukhanova [2023] and *σ_k_* is the standard error for the expected pup proportion for species *k* on the logit scale (we we set such that the implied coefficient of variation was 0.2). Note that by shrinking regression coefficients towards zero, our prior on ***β*** induces regularization [Hobbs and Hooten, 2015]; the particular form of the variance-covariance matrix also puts regression coefficients on the same footing, regardless of covariate scale.

Models with prior distributions for misclassification and thinning parameters were either numerically unstable or resulted in posterior estimates that differed substantially from prior means. This behavior is undesirable since there is presumably little information in count data alone about detection or misclassification. Such a shift is likely the result of model misspecification or confounding between detection and density processes [Bravington et al., 2021]. Instead, we relied on a parametric bootstrap procedure to propagate uncertainty in misclassification and detection processes into resultant estimates [for a detailed description of the bootstrapping procedure, see Conn and Ferguson, 2026]. Point estimates of abundance and density-habitat parameters were conditioned on point estimates of detection and misclassification parameters.

We fitted our models using maximum marginal likelihood procedures implemented in Template Model Builder (TMB) software [Kristensen et al., 2015]. Under this framework, parameters with prior distributions (in our case, the logits of pup proportions and covariate regression parameters) were treated as random effects and integrated out of the joint likelihood using Laplace approximation. We used parametric bootstrap estimates of uncertainty to construct 95% log-based confidence intervals for abundance [Burnham et al., 1987, Buckland et al., 2001]. Code and data to recreate our analyses are available at https://github.com/pconn/BOSSrussia/ and will be archived to a publicly available data repository upon manuscript acceptance.

### Goodness-of-fit

We used probability integral transforms [Conn et al., 2018] in the form of randomized quantile residuals [Dunn and Smyth, 1996] to detect possible lack-of-fit in count data. The specific algorithm used is provided in Boveng et al. (2025). For each species, we also recorded several metrics to help diagnose whether models included potentially problematic extrapolations (i.e., predictions where combinations of covariates produced predictions that are much higher than in sampled cells). For each species *k*, let *µ_k,max_* give the maximum prediction over space-time indices that were actually sampled. Then let *n_k_* = Σ*_j_ I_µk,j>µk,max_* represent the total number of predictions that exceed the maximum observed value. Additionally, as a measure of the influence of such extrapolations, we computed Λ*_k_* = max*_j_*(*µ_k,j_/µ_k,max_*), which measures how much larger the most extreme prediction is relative the largest prediction in a surveyed cell.

### Sensitivity analyses

In addition to our preferred models, we fit reduced parameter versions where each predictive covariate was sequentially dropped from the model structure to investigate the influence of model specification on resultant estimates. We also fitted alternative models that omitted species misclassification and used different assumptions about ringed seal availability and lair use. Information about sensitivity runs is provided in *Supplementary Information 1*.

## Results

Over both years (2012 and 2013), surveyors detected 12,624 seals in the wBS, of which 5,748 (46%) were photographed. Photographed seals included 655 bearded seals, 979 ribbon seals, 989 ringed seals, 2,437 spotted seals, 457 seals of unknown species, and 231 pups. In the SO, researchers detected 5,730 seals in the surveyed area, of which 4,360 (76%) were photographed. These included 453 bearded seals, 1805 ribbon seals, 844 ringed seals, 721 spotted seals, 102 seals of unknown species, and 435 pups.

In the wBS, we estimated that abundance of bearded, ribbon, ringed, and spotted seals from our preferred model was 185,330, 68,564, 272,609, and 226,003, respectively in 2012 and 144,105, 39,435, 203,130 and 90,501 in 2013 (Table 2). For reference, mean haul-out (i.e., availability) probabilities (range) for these species used in our models were 0.29 (0.11-0.45), 0.47 (0.23-0.72), 0.27 (0.07-0.34), and 0.51 (0.35-0.64) in 2012 and 0.25 (0.11-0.48), 0.35 (0.15-0.67), 0.24 (0.05-0.34), and 0.45 (0.30-0.62) in 2013, respectively. Bearded and ringed seals were concentrated in the northern third of the study area, whereas ribbon and spotted seals were concentrated on the southern edges of pack ice; densities of all four species were also high in and around Karaginsky Bay (Figs. 3-4). Ribbon and spotted seal distributions appeared constricted in 2013 when compared to 2012, with ribbon seal densities concentrated at the southern ice edge, and spotted seal densities concentrated in Karaginsky Bay.

**Figure 3:**
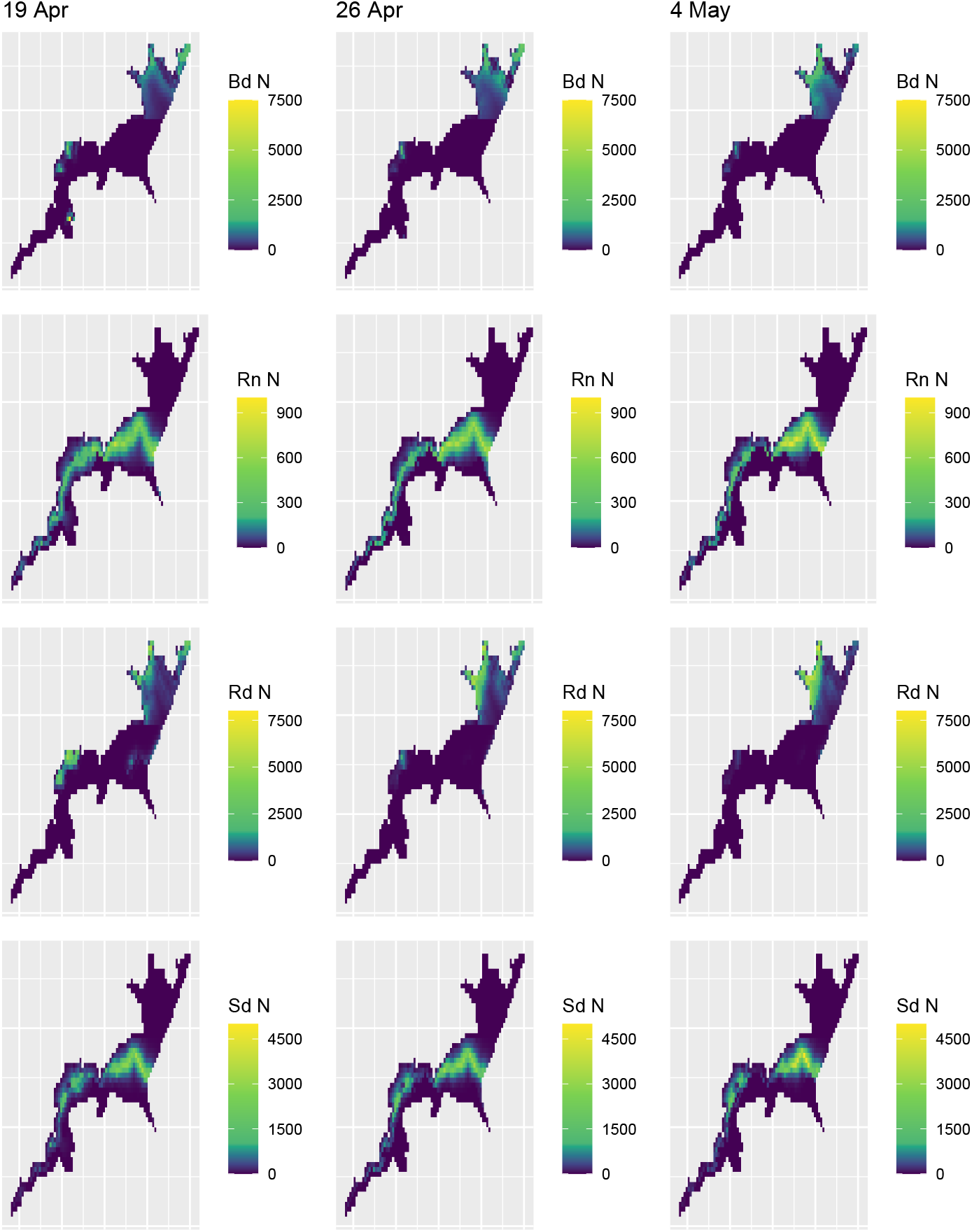
Estimates of bearded (Bd), ribbon (Rn), ringed (Rd), and spotted (Sd) seal abundance on three representative days during the spring of 2012 in the western Bering Sea, as predicted by a statistical model fitted to aerial survey counts.

**Figure 4:**
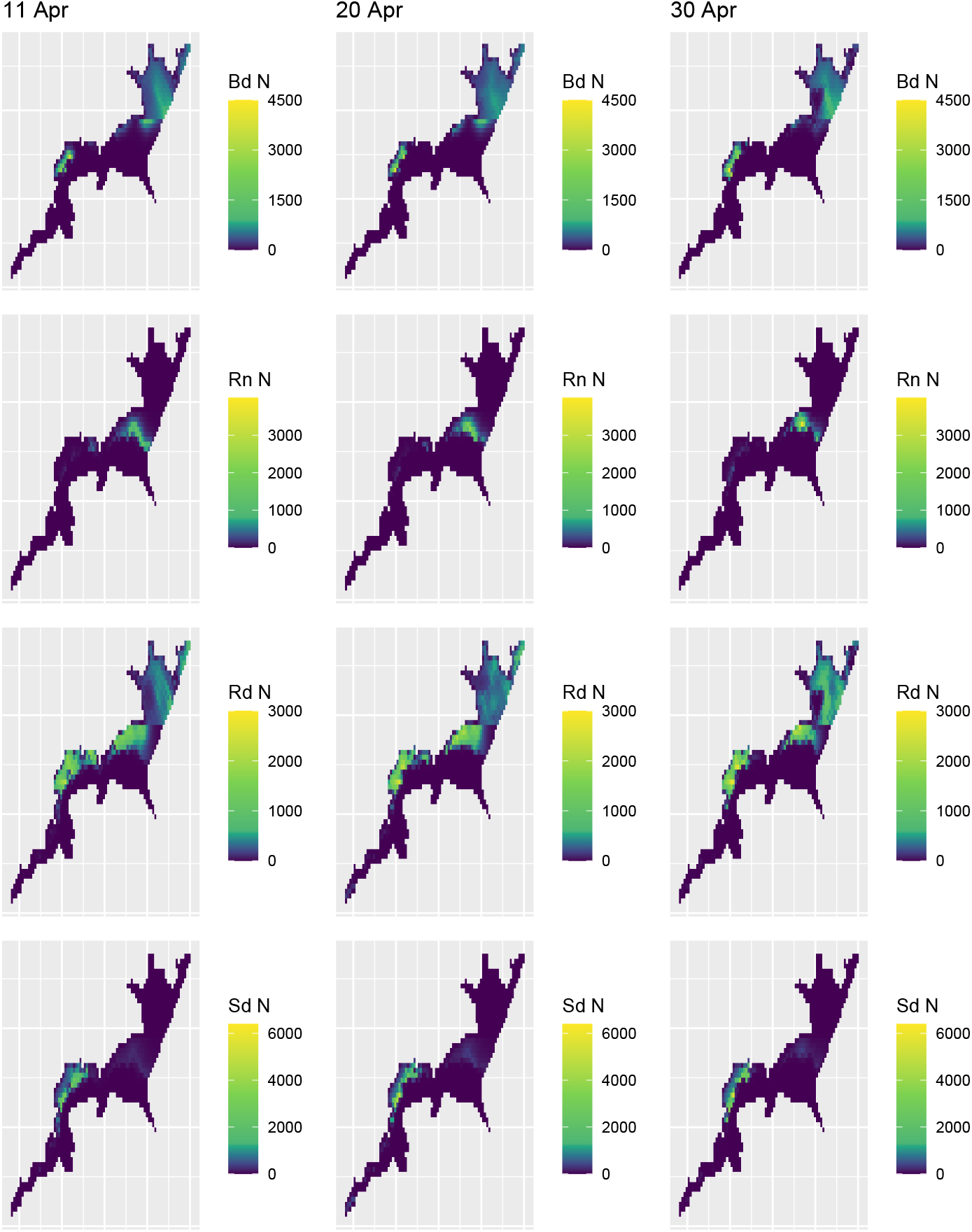
Estimates of bearded (Bd), ribbon (Rn), ringed (Rd), and spotted (Sd) seal abundance during three representative days during the spring of 2013 in the western Bering Sea, as predicted by a statistical model fitted to aerial survey counts. Note that the scales differ from 2012 plots to better highlight areas of high and low abundance.

**Table 2:**
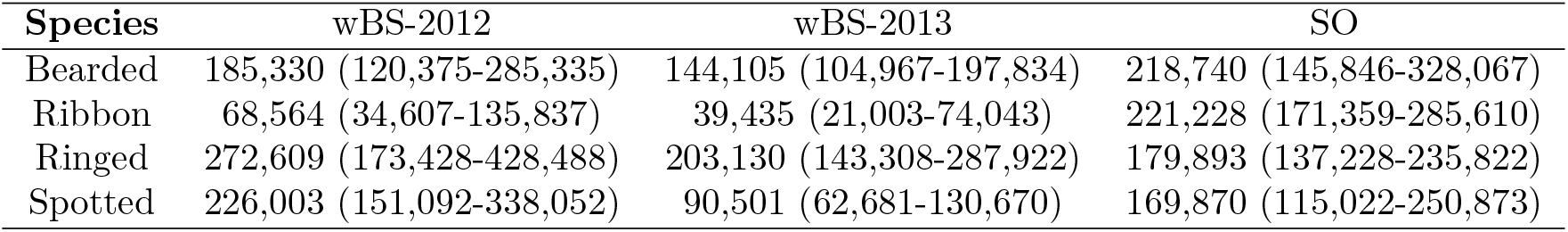
Estimates of ice seal abundance in the western Bering Sea and Sea of Okhotsk in 2012 and 2013, together with 95% confidence intervals.

| Species | wBS-2012 | wBS-2013 | SO |
| --- | --- | --- | --- |
| Bearded | 185,330 (120,375-285,335) | 144,105 (104,967-197,834) | 218,740 (145,846-328,067) |
| Ribbon | 68,564 (34,607-135,837) | 39,435 (21,003-74,043) | 221,228 (171,359-285,610) |
| Ringed | 272,609 (173,428-428,488) | 203,130 (143,308-287,922) | 179,893 (137,228-235,822) |
| Spotted | 226,003 (151,092-338,052) | 90,501 (62,681-130,670) | 169,870 (115,022-250,873) |

In 2012, spotted seals seemed to be most prone to extrapolation bias, with *n* = 252 predictions (15.8/day) exceeding the maximum prediction for sampled cells; the maximum prediction in unsampled cells was Λ*_k_* = 2.1 times greater than that for sampled cells (see Online Resource 2D). By contrast, bearded, ribbon, and ringed seals had *n* = 33 (2.1/day), *n* = 5 (0.3 cells/day), and *n* = 89 (5.6 cells/day) predictions in unsampled cells exceed the maximum for sampled cells. Maximum predictions were Λ*_k_* = 3.0, 1.0, and 1.5 times greater in unsampled cells than in sampled cells for these species (*Supplementary Information 1*).

In 2013, extrapolation metrics suggested that just *n* = 1 bearded seal prediction was an extrapolation, compared to *n* = 120 (6 cells/day) for ribbon seals, *n* = 35 (1.8/day) for ringed seals, and *n* = 89 (4.4/day) for spotted seals (*Supplementary Information 1*). The maximum prediction in unsampled cells for these species was Λ*_k_* = 1.0, 2.7, 1.2, and 2.0 times greater than for sampled cells, respectively. Large ribbon seal predictions were notable in the southwest corner of sea ice that was never sampled (compare Figs. 1 & 4).

In the Sea of Okhotsk, estimates of bearded, ribbon, ringed, and spotted seal abundance from our preferred models were 218,740, 221,228, 179,893, and 169,870, respectively (Table 2). For reference, the mean values for availability for each species (range) used in the SO analysis were 0.24 (0.16-0.27), 0.54 (0.33-0.62), 0.44 (0.24-0.51), and 0.57 (0.43-0.61), respectively. Spotted and ribbon seals were common along the ice edge, with ribbon seals having highest densities in the southwest and spotted seals having highest densities in the northeast. Bearded and ringed seals were spread more evenly including the interior of the pack ice, though bearded seal densities appeared highest in nearshore areas while ringed seal densities were highest offshore (Fig. 5, *Supplementary Information 2*). Our model predicted relatively high numbers of bearded and spotted seals in the northeast of the Okhotsk Sea, an area which was not surveyed. Predictions in this area should be treated with caution. Goodness-of-fit tests suggested some lack-of-fit, particularly for spotted, ribbon, and unknown seal counts (*Supplementary Information 2*). However, extrapolation metrics suggested that extrapolation in unsurveyed locations was not a large cause of bias (*Supplementary Information 1*).

**Figure 5:**
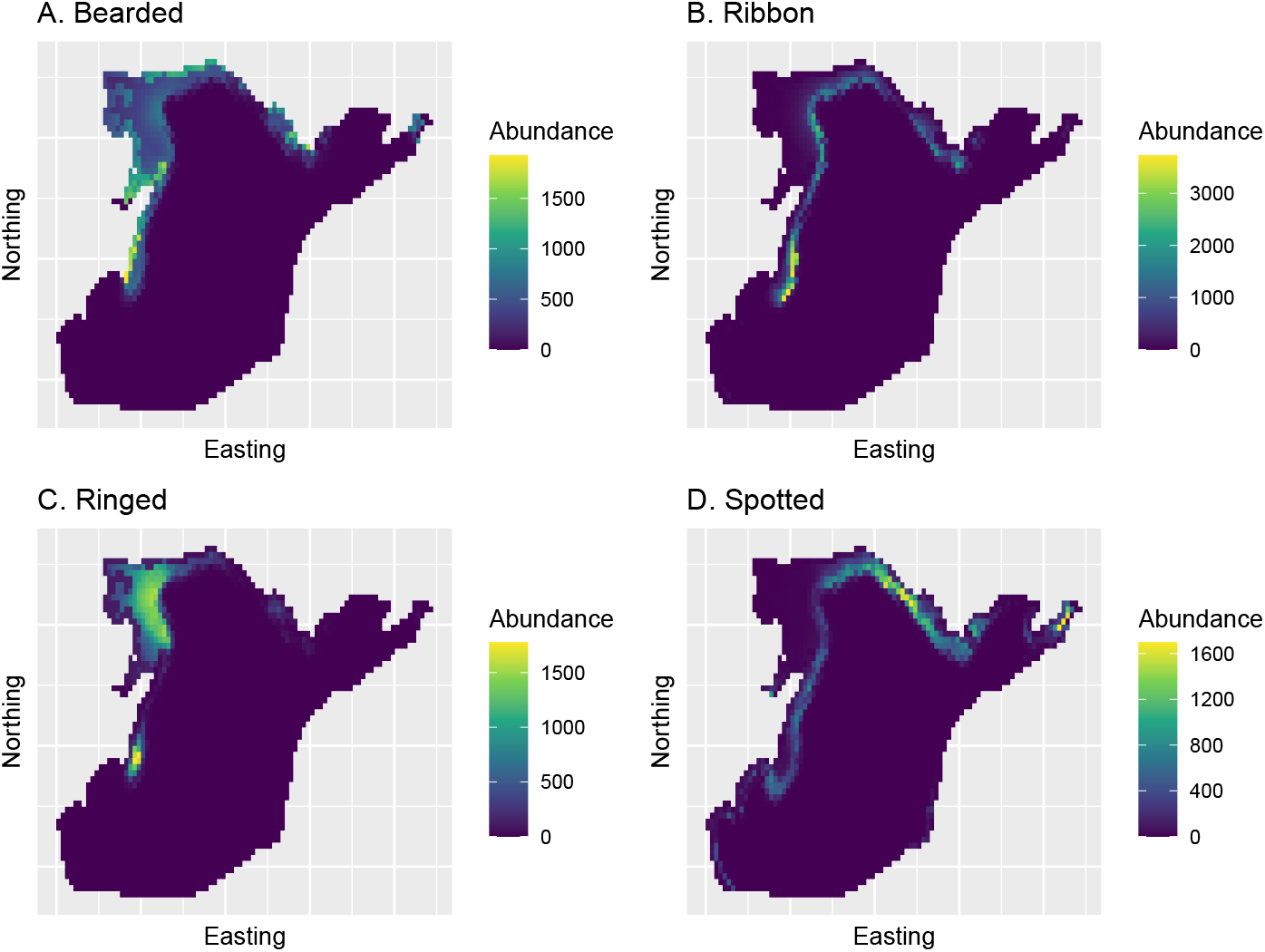
Estimates of bearded, ribbon, ringed, and spotted seal abundance during early May of 2013 in the Sea of Okhotsk, as predicted by a statistical model fitted to aerial survey counts.

Sensitivity runs for the Sea of Okhotsk indicated considerable differences in estimates depending upon model assumptions (*Supplementary Information 1*). For instance, removal of species misidentification resulted in markedly different estimates of abundance for bearded, ringed, and spotted seals. Similarly, an alternative model using ringed seal availability predictions from the Bering Sea resulted in much higher estimates of ringed seal abundance.

## Discussion

In this paper, we reanalyzed Russian aerial survey data to estimate population size and spatial distribution of ice-associated seals in the western Bering Sea (2012 and 2013) and the Sea of Okhotsk (2013 only). Importantly, we accounted for a number of nuisance detection processes that were largely unaddressed in initial publications [Chernook et al., 2014, 2018]. These modifications increased abundance estimates because we adjusted for seals that were in the water and thus unavailable to be counted; however, differences in availability combined with various species identification errors meant that the amount of increase for each species varied considerably. For instance, in the Sea of Okhotsk, the smallest increase was for ribbon seals (22%), while the largest was for bearded seals (445%).

Although we did the best we could with the data at hand, there were considerable issues that had to be overcome, including nuisance detection processes, incomplete spatial coverage during surveys, and changing sea ice habitat while surveys were being conducted (especially in the western Bering). Gaps in spatial coverage were largely a function of the large areas being surveyed, coupled with challenging logistical constraints when surveying the Russian Far East and a limited budget for survey efforts. As a consequence, models fitted to survey data could be sensitive to the covariates used to predict seal abundance in unsampled locations; particularly for ribbon and spotted seals who preferred the southern edge of pack ice in the western Bering that was difficult to access. We thus relied on relatively simple predictive covariate functional forms in order to reduce extrapolative error that could positively bias abundance estimates.

Although the subject of extrapolation has been addressed in the species distribution literature [e.g. Conn et al., 2015a, Mannocci et al., 2015], sometimes under the label of “transferability” [Randin et al., 2006], there are no clear answers on how to best balance prediction metrics often used for model selection (e.g., AIC, cross validation) with more subjective indicators of extrapolation (such as at the unsampled edge of a study area). Our approach in this paper was to limit extrapolation by simplifying our models until we arrived at a structure that seemed to avoid excessive extrapolation. We suspect this will continue to be an area of “uncharted waters” [Sequeira et al., 2018] when complicated spatial or spatio-temporal models are fitted to marine mammal counts.

Our estimates of uncertainty (e.g., standard errors, confidence intervals) in the present study are likely understated. For instance, since we were restricted to simple formulations for habitat covariates, we are not accounting for spatial autocorrelation that is common in ecological data [Lichstein et al., 2002]. Accounting for spatial autocorrelation would tend to decrease precision. One indication that uncertainty is underestimated is the lack-of-fit observed for some of the survey counts (*Supplementary Information 2*). A usual approach in these situations is to calculate a variance inflation factor [Burnham and Anderson, 2002], but it is unclear how this should best be done for a multi-species count model using different types of observations. Similarly, the differences among sensitivity runs point to additional model uncertainty in estimates that is not addressed with single-model confidence intervals. However, the level of model curation needed to avoid extrapolation bias does not fit well with common model averaging paradigms [e.g., Burnham and Anderson, 2002], so our preference is to simply acknowledge that our confidence intervals are likely too narrow.

A possible source of bias in our estimates has to do with availability probability. In particular, we based availability probability on long term average haul-out behavior of seals with bio-loggers deployed in the Bering, Chukchi, and Beaufort Seas [Boveng et al., 2026, London et al., 2024]. If seals hauled out on ice at different rates in 2012 and 2013 than their long term averages, our estimates may be biased to some degree. Similarly, if seal haul-out behavior differed between surveyed locations and locations where bio-loggers were deployed, abundance estimates there could be biased. Finally, ringed seal predictions were made based on behavior observed in the Chukchi Sea and did not include a correction for seals that may have remained hidden in subnivean lairs. For these reasons, ringed seal availability corrections are probably the least reliable and abundance estimates are likely biased low in the wBS if some ringed seals construct snow lairs there.

Subject to the above caveats, there appeared to be close to 200,000 of each species (bearded, ribbon, ringed, and spotted seals) in 2013 in the Sea of Okhotsk. Point estimates in the western Bering Sea declined between 2012 and 2013 for all four species: from 185,000 to 145,000 (−22%) for bearded seals; 69,000 to 39,000 (−43%) for ribbon seals; 270,000 to 203,000 (−25%) for ringed seals; 226,000 to 91,000 (−60%) for spotted seals), though only spotted seals had confidence intervals that did not overlap. These apparent declines mirror similar declines in the eastern Bering Sea in the same years, where percent declines were −7%, −64%, −27%, and −54%, respectively [Boveng et al., 2026]. This suggests that a similar dynamic occurred in Russian and U.S. waters of the Bering Sea between 2012 and 2013. However, we are unable to determine the underlying cause. For instance, it could have been an actual population decline following an unusual mortality event (UME) that involved skin lesions and other symptoms [NOAA, 2012, 2018], or it could have been due to a dramatic shift in haul-out behavior of these species compared to their long-term averages.

Implementing surveys across political boundaries is logistically and politically difficult. Yet, it is the only way to produce comprehensive estimates of transboundary, ecologically relevant units such as distinct population segments (DPSs). For example, by combining the mean estimate of bearded seals from this paper with those from the eastern Bering [Boveng et al., 2026] and Chukchi [Boveng et al., 2025] Seas conducted at the same time of year, we arrive at a Beringia subpopulation of *N*^^^*_bd_* = 573, 000. Similarly, we can combine ribbon and spotted seal estimates from the Bering Sea and Sea of Okhotsk to arrive at something close to a range-wide estimate for these species, though these numbers vary considerably based on which year is chosen for Bering Sea estimates (i.e., 2012 vs 2013). Conservatively, using 2013 (when lower seal abundance was estimated in the Bering Sea) provides estimates of 286,000 ribbon seals and 395,000 spotted seals [not including approximately 5000 spotted seals in the Sea of Japan and Yellow Sea; Trukhin, 2024, Yan et al., 2018]. To our knowledge, these are the first survey-based, range-wide estimates for these species. In addition to population estimates, our models provide spatially-explicit predictions of seal densities. These maps should be useful for marine planning and for monitoring how seal distributions respond to changing sea ice conditions.

Future surveys will be needed to establish long term trends in seal abundance. This is especially important given widespread conservation concern for seals that rely on spring sea ice for critical life history functions. For instance, the Beringia DPS of bearded seals and the Arctic and Okhotsk subspecies of ringed seals are currently listed as threatened under the U.S. Endangered Species Act [National Marine Fisheries Service, 2012b,a], largely because of the likelihood of future habitat declines. Future surveys could be improved by (1) increasing spatial and temporal replication to reduce extrapolation bias, (2) simultaneously deploying satellite-linked bio-loggers to better estimate availability probability in surveyed areas, and (3) employing double sampling and behavioral response trials to estimate other nuisance detection parameters such as sensor perception probability, species misclassification probabilities, and seal response to aircraft.

## Acknowledgements

We thank A. Warlick and S. Dahle for comments on an earlier draft that improved its quality. Funding for aerial surveys was provided by the U.S. National Oceanic and Atmospheric Administration, the U.S. Bureau of Ocean Energy Management, and the Marine Mammal Council of Russia. Views expressed are those of the authors and do not necessarily represent findings or policy of any government agency. Use of trade or brand names does not indicate endorsement by the U.S. government.

## Author contributions statement

P.B. and V.C. conceived of the surveys; A.V. and V.C. planned the surveys; P.C. analyzed the data and drafted the manuscript; I.T. conducted preliminary analysis of Russian Federation survey data and provided translations and interpretation of data. All authors edited the manuscript.

## Supplementary Information 1

### Description of sensitivity analyses and extrapolation metrics for seal abundance models

We ran a number of sensitivity analyses to investigate how abundance estimates were impacted by alternate combinations of predictive covariates. For both the Sea of Okhotsk (SO) and western Bering Sea (wBS), we sequentially removed candidate predictors from each seal species abundance intensity model; in a few cases, we also added predictors to show what could happen with extrapolation bias. We next describe sensitivity analyses for the wBS and SO.

### wBS sensitivity analysis

In the wBS, we ran a similar set of sensitivity analyses, but in this case removed habitat predictors one at a time for each species owing to greater overall sensitivity of results to model assumptions. One major differences between the two analysis approaches was that bearded and ringed seals permitted more highly parameterized models than for ribbon and spotted seals. Ribbon and spotted seals were problematic in the wBS in that densities were often highest towards the eastern ice edge and transects did not always extend all the way out to these areas. This often led to highly parameterized models predicting unrealistically high numbers of seals in locations that weren’t sampled. Our preferred models for ribbon and spotted seals were thus relatively simple.

For bearded and ringed seals, our preferred model related latent abundance (*Z_kj_*) for each species *k* to covariates in cell *j* using a loglinear modeling framework. Using the “formula” notation common to the R programming environment linear modeling functions (e.g., lm, glm; R Development Core Team, 2017), we write a model for log(*µ_kj_*) in 2012 as

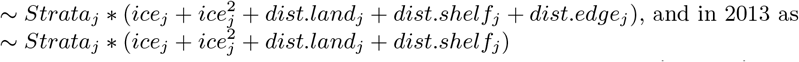

That is, there were strata-specific effects of covariates representing sea ice concentration (*ice_j_*) and distance from various features (dist.land: distance from land; dist.shelf: distance from 1000 m isobath; dist.edge: distance from ice edge). The models differed between years because inclusion of *dist.edge* resulted in clear extrapolations (see below).

For ribbon and spotted seals, our preferred model for log(*Z_kj_*) in both years was *~ Strata_j_ + ice_j_ + ice^2^_j_ + dist.shelf_j_*.

In this case, spatial strata was a simple additive effect, reducing the number of parameters.

For each species and year, we sequentially left out terms for each species and recorded the impact on the fitted log likelihood, the abundance estimate of each species, and on extrapolation metrics defined in the main text. We also investigated effects of adding covariates (*dist.land* and *dist.edge*, for years and species for which they were missing). In 2012 in the wBS, abundance estimates of all species were fairly sensitive to model structure (Table S1.1). However, bearded seal models with the highest log likelihoods (those best supported by the data) all seemed to agree on estimates in the 180,000-185,000 range. Ribbon seal models varied widely, including estimates from 44,000-94,000, although the model with 94,000 (adding *dist.land*) was also accompanied with increased evidence of extrapolation in unsampled areas. Thus, an estimate in the 44,000-65,000 range seems most appropriate. Ringed seal estimates were mostly near 270,000, though some runs (e.g., omitting spatial strata) resulted in a higher estimate. As one example of clear extrapolation (Table S1.2), substracting *dist.land* from the ringed seal model resulted in an estimate close to 3,000,000; in this case, there were 841 additional extrapolations compared to the main model, with at least one unsampled cell predicted to have an abundance that was 155 times higher that in any sampled cell. Spotted seal estimates ranged from 180,000 to 237,000 for models with high log likelihoods, with no clear model performing the best. Estimates for individual species were often impacted by changes to model structure of other species, which is to be expected given how species misidentification was modeled.

In 2013, abundance estimates were similarly sensitive to model structure, although to a lesser degree (Table S1.3). For bearded seals, estimates of highly supported models were betweeen 144,000 and 162,000, with most on in the 140,000-150,000 range. For ribbon seals, highly supported models produced estimates between 22,000 and 45,000; the model adding *dist.edge* was a conspicuous example of extrapolation, with an estimate near 358,000 and anomalously high predictions in unsampled cells near the ice edge (Table S1.4). Highly ranked ringed seal models produced estimates near 200,000 (at least, when discarding the model that added *dist.edge* which also exhibited unfavorable extrapolation metrics). Finally, highly ranked spotted seal models produced estimates in the 90,000-105,000 range.

### SO sensitivity analysis

In the SO, we used a base model for each species where habitat preferences included (1) linear and quadratic effects for sea ice concentration, (2) linear and quadratic effects of *depth*^1*/*3^, and linear effects for distance from land, distance from shelf break, and distance from ice edge. We arrived at this model after experimenting with other configurations and striving for a model which maintained as much flexibility as possible while avoiding potentially problematic extrapolations to areas that weren’t well sampled. Our base model used predictions of ringed seal availability based on seals that were tagged in the Chukchi Sea (London et al., 2024). After fitting this “base model,” we fit alternative models where environmental covariates were removed one at a time. We also fit a model where ringed seal availability was based on seals in the Bering Sea (this sample is biased towards younger animals, who spend less time hauled-out), as well as model where species misclassification rates were set to 0.0.

Models fit to seal counts in the SO were somewhat sensitive to model structure (Table S1.5). This was particularly the case for ringed seals when alternative availability schedules based on Bering Sea seals were employed (Table S1.5). The impact of ignoring species misidentification was also profound, with markedly different abundance estimates if this source of possible bias wasn’t addressed. Fortunately, we identified relatively few, low magnitude extrapolations (defined as predictions in unsampled grid cells being greater than those in sampled grid cells; see Table S1.6). Owing to the large difference in AIC score for our base model (Table S1.5), we suggest these estimates are likely to most appropriate for management (i.e., roughly 220,000 bearded and ribbon seals, 180,000 ribbon seals, and 170,000 spotted seals).

---

**Table S1.1:** Results of sensitivity runs for 2012 in the wBS. The first column gives sensitivity run name, where “main” is the preferred model presented in the main paper, “no misID” is an attempt at running the model without species misidentification (this model did not converge so no results are reported), and “rd Bering” use Bering Sea predictions of ringed seal availability. For the remaining models, ‘-’ and ‘+’ indicate whether a particular covariate is added or subtracted from the main model, and (sp) gives the species for which a covariate was added or substracted (bd: bearded; rn: ribbon; rd: ringed; sd: spotted). For each sensitivity run, we present the optimized log likelihood (LogL), the number of regression parameters (*k*), the difference in AIC score for each model compared to the “main” model, and abundance estimates for each species (Nbd - Nsd). The AIC score only included the number of regression parameters in the parameter count; positive values indicate increased predictive performance relative to our preferred model.

| Model | LogL | k | ΔAIC | $N_{bd}$ | $N_{rn}$ | $N_{rd}$ | $N_{sd}$ |
| --- | --- | --- | --- | --- | --- | --- | --- |
| main | -1633.91 | 44 | 0.00 | 185330 | 68564 | 272609 | 226003 |
| no misID | NA | 44 | NA | NA | NA | NA | NA |
| rd Bering | -1634.98 | 44 | -2.14 | 182512 | 68440 | 176335 | 230386 |
| -strata (bd) | -1651.72 | 35 | -17.63 | 220612 | 68544 | 264262 | 223535 |
| -dist_land (bd) | -1642.63 | 42 | -13.44 | 166059 | 68919 | 293312 | 225291 |
| -dist_shelf (bd) | -1634.52 | 42 | 2.78 | 185050 | 68712 | 274692 | 226454 |
| -dist_edge (bd) | -1638.12 | 42 | -4.42 | 183229 | 68300 | 268545 | 224807 |
| -strata (rn) | -1633.91 | 42 | 4.00 | 185125 | 44395 | 272541 | 246628 |
| -dist_shelf (rn) | -1635.40 | 43 | -0.98 | 186244 | 65954 | 272857 | 235788 |
| +dist_edge (rn) | -1639.31 | 45 | -12.80 | 184757 | 65778 | 271990 | 236285 |
| +dist_land (rn) | -1631.79 | 45 | 2.24 | 185166 | 93871 | 272529 | 223717 |
| -strata (rd) | -1651.05 | 35 | -16.28 | 180693 | 67542 | 311852 | 233532 |
| -dist_land (rd) | -1640.12 | 42 | -8.42 | 183944 | 67269 | 3012168 | 223130 |
| -dist_shelf (rd) | -1634.20 | 42 | 3.42 | 184591 | 68933 | 279506 | 229750 |
| -dist_edge (rd) | -1639.73 | 42 | -7.65 | 183433 | 69216 | 268192 | 233725 |
| -strata (sd) | -1634.62 | 42 | 2.58 | 185231 | 77707 | 272551 | 190646 |
| -dist_shelf (sd) | -1659.48 | 43 | -49.14 | 189241 | 93196 | 266579 | 184323 |
| +dist_edge (sd) | -1633.15 | 45 | -0.48 | 185306 | 68764 | 272126 | 236785 |
| +dist_land (sd) | -1633.91 | 45 | -2.00 | 185358 | 68567 | 272609 | 224906 |

**Table S1.2:** Extrapolation metrics associated with sensitivity runs for 2012 in the wBS. The first column gives sensitivity run name, where “main” is the preferred model presented in the main paper, “no misID” is an attempt at running the model without species misidentification (this model did not converge so no results are reported), and “rd Bering” used Bering Sea predictions of ringed seal availability. For the remaining models, ‘-’ and ‘+’ indicate whether a particular covariate is added or subtracted from the main model, and (sp) gives the species for which a covariate was added or substracted (bd: bearded; rn: ribbon; rd: ringed; sd: spotted). For each sensitivity run, we present the number of spatio-temporal cells for which abundance of a given species was predicted to be greater than the maximum predicted for sampled cells *n_species_*, as well as the ratio of the maximum prediction in unsampled cells to the maximum prediction in sampled cells (Λ*_species_*).

| Model | $n_{bd}$ | $n_{rn}$ | $n_{rd}$ | $n_{sd}$ | $\Lambda_{bd}$ | $\Lambda_{rn}$ | $\Lambda_{rd}$ | $\Lambda_{sd}$ |
| --- | --- | --- | --- | --- | --- | --- | --- | --- |
| main | 33 | 5 | 89 | 252 | 2.96 | 1.04 | 1.52 | 2.10 |
| no misID | NA | NA | NA | NA | NA | NA | NA | NA |
| rd Bering | 11 | 5 | 139 | 251 | 4.52 | 1.04 | 2.11 | 2.14 |
| -strata (bd) | 33 | 5 | 89 | 101 | 2.80 | 1.04 | 1.52 | 1.54 |
| -dist_land (bd) | 72 | 180 | 89 | 7 | 4.38 | 1.83 | 1.53 | 1.01 |
| -dist_shelf (bd) | 32 | 5 | 88 | 197 | 2.79 | 1.05 | 1.52 | 2.08 |
| -dist_edge (bd) | 33 | 5 | 89 | 248 | 2.99 | 1.04 | 1.52 | 2.08 |
| -strata (rn) | 33 | 3 | 89 | 385 | 2.98 | 1.04 | 1.52 | 2.41 |
| -dist_shelf (rn) | 37 | 51 | 89 | 259 | 3.32 | 1.07 | 1.52 | 2.25 |
| +dist_edge (rn) | 33 | 36 | 89 | 321 | 2.89 | 1.53 | 1.52 | 2.19 |
| +dist_land (rn) | 33 | 423 | 89 | 244 | 2.88 | 2.28 | 1.52 | 2.04 |
| -strata (rd) | 153 | 5 | 60 | 248 | 1.81 | 1.04 | 2.30 | 2.07 |
| -dist_land (rd) | 246 | 5 | 92 | 246 | 2.69 | 1.04 | 2.17 | 2.05 |
| -dist_shelf (rd) | 70 | 5 | 92 | 252 | 1.61 | 1.04 | 1.53 | 2.10 |
| -dist_edge (rd) | 16 | 4 | 76 | 247 | 2.02 | 1.04 | 1.48 | 2.05 |
| -strata (sd) | 20 | 8 | 182 | 242 | 2.41 | 1.07 | 2.10 | 2.07 |
| -dist_shelf (sd) | 17 | 5 | 931 | 251 | 2.18 | 1.04 | 156.66 | 2.02 |
| +dist_edge (sd) | 66 | 5 | 120 | 257 | 2.01 | 1.04 | 1.74 | 2.15 |
| +dist_land (sd) | 79 | 7 | 109 | 255 | 1.81 | 1.05 | 1.52 | 2.17 |

**Table S1.3:** Results of sensitivity runs for 2013 in the wBS. The first column gives sensitivity run name, where “main” is the preferred model presented in the main paper, “no misID” is a model without species misidentification, and “rd Bering” used Bering Sea predictions of ringed seal availability. For the remaining models, ‘-’ and ‘+’ indicate whether a particular covariate is added or subtracted from the main model, and (sp) gives the species for which a covariate was added or substracted (bd: bearded; rn: ribbon; rd: ringed; sd: spotted). For each sensitivity run, we present the optimized log likelihood (LogL), the number of regression parameters (*k*), the difference in AIC score for each model compared to the “main” model, and abundance estimates for each species (Nbd - Nsd). The AIC score only included the number of regression parameters in the parameter count; positive values indicate increased predictive performance relative to our preferred model.

| Model | LogL | k | ΔAIC | $N_{bd}$ | $N_{rn}$ | $N_{rd}$ | $N_{sd}$ |
| --- | --- | --- | --- | --- | --- | --- | --- |
| main | -1926.46 | 38 | 0.00 | 144105 | 39435 | 203130 | 90501 |
| no misID | -1912.99 | 38 | 26.94 | 138197 | 44133 | 227465 | 80079 |
| rd Bering | -1942.09 | 38 | -31.24 | 150409 | 39184 | 420815 | 99194 |
| -strata (bd) | -1955.82 | 29 | -40.71 | 166127 | 41495 | 217272 | 95460 |
| -dist_land (bd) | -1941.64 | 36 | -26.35 | 144967 | 37669 | 215152 | 93357 |
| -dist_shelf (bd) | -1929.20 | 36 | -1.47 | 161824 | 38412 | 205441 | 90567 |
| +dist_edge (bd) | -1926.15 | 36 | 4.63 | 140869 | 39502 | 203179 | 90068 |
| -strata (rn) | -1926.41 | 36 | 4.11 | 144643 | 22131 | 205321 | 89680 |
| -dist_shelf (rn) | -1934.86 | 37 | -14.79 | 144449 | 27598 | 203704 | 90765 |
| +dist_edge (rn) | -1938.16 | 39 | -25.40 | 144351 | 357766 | 205949 | 91731 |
| +dist_land (rn) | -1924.06 | 39 | 2.81 | 143943 | 36764 | 202909 | 90567 |
| -strata (rd) | -1962.45 | 29 | -53.97 | 148011 | 42380 | 164027 | 107311 |
| -dist_land (rd) | -1935.94 | 36 | -14.96 | 149327 | 40148 | 185105 | 93390 |
| -dist_shelf (rd) | -1932.50 | 36 | -8.08 | 143975 | 40158 | 198466 | 91398 |
| -dist_edge (rd) | -1910.02 | 36 | 36.90 | 141903 | 39065 | 242274 | 109782 |
| -strata (sd) | -1932.48 | 36 | -8.04 | 152071 | 32899 | 226811 | 165388 |
| -dist_shelf (sd) | -1932.84 | 37 | -10.75 | 145412 | 41755 | 205026 | 83635 |
| +dist_edge (sd) | -1923.38 | 39 | 4.16 | 147970 | 45122 | 209494 | 104827 |
| +dist_land (sd) | -1926.41 | 39 | -1.89 | 144183 | 39438 | 203380 | 89657 |

**Table S1.4:** Extrapolation metrics associated with sensitivity runs for 2013 in the wBS. The first column gives sensitivity run name, where “main” is the preferred model presented in the main paper, “no misID” is a model without species misidentification, and “rd Bering” used alternative predictions of ringed seal availability. For the remaining models, ‘-’ and ‘+’ indicate whether a particular covariate is added or subtracted from the main model, and (sp) gives the species for which a covariate was added or substracted (bd: bearded; rn: ribbon; rd: ringed; sd: spotted). For each sensitivity run, we present the number of spatio-temporal cells for which abundance of a given species was predicted to be greater than the maximum predicted for sampled cells *n_species_*, as well as the ratio of the maximum prediction in unsampled celsl to the maximum prediction in sampled cells (Λ*_species_*).

| Model | $n_{bd}$ | $n_{rn}$ | $n_{rd}$ | $n_{sd}$ | $\Lambda_{bd}$ | $\Lambda_{rn}$ | $\Lambda_{rd}$ | $\Lambda_{sd}$ |
| --- | --- | --- | --- | --- | --- | --- | --- | --- |
| main | 1 | 120 | 35 | 89 | 1.01 | 2.66 | 1.23 | 2.05 |
| no misID | 0 | 138 | 46 | 60 | 1.00 | 2.20 | 1.33 | 1.93 |
| rd Bering | 1 | 120 | 210 | 91 | 1.00 | 2.67 | 1.71 | 1.98 |
| -strata (bd) | 1 | 126 | 111 | 88 | 1.01 | 2.58 | 14.96 | 1.76 |
| -dist_land (bd) | 2 | 120 | 35 | 8 | 1.01 | 2.73 | 1.30 | 1.03 |
| -dist_shelf (bd) | 2 | 120 | 26 | 30 | 1.01 | 2.79 | 1.19 | 1.79 |
| +dist_edge (bd) | 1 | 120 | 35 | 94 | 1.01 | 2.66 | 1.24 | 1.96 |
| -strata (rn) | 1 | 186 | 33 | 81 | 1.01 | 2.74 | 1.22 | 1.99 |
| -dist_shelf (rn) | 2 | 387 | 33 | 83 | 1.01 | 1.30 | 1.23 | 2.02 |
| +dist_edge (rn) | 1 | 357 | 38 | 95 | 1.01 | 21.92 | 1.25 | 2.07 |
| +dist_land (rn) | 1 | 84 | 36 | 89 | 1.01 | 2.37 | 1.23 | 2.05 |
| -strata (rd) | 185 | 120 | 46 | 62 | 1.84 | 2.74 | 1.31 | 1.83 |
| -dist_land (rd) | 4 | 121 | 42 | 84 | 1.35 | 2.69 | 1.33 | 1.97 |
| -dist_shelf (rd) | 0 | 121 | 35 | 98 | 1.00 | 2.67 | 1.24 | 2.07 |
| -dist_edge (rd) | 5 | 120 | 37 | 91 | 1.06 | 2.65 | 1.22 | 2.06 |
| -strata (sd) | 3 | 119 | 6 | 94 | 1.07 | 2.63 | 1.08 | 2.00 |
| -dist_shelf (sd) | 3 | 119 | 6 | 98 | 1.02 | 2.64 | 1.05 | 2.12 |
| +dist_edge (sd) | 2 | 120 | 35 | 89 | 1.01 | 2.69 | 1.24 | 2.03 |
| +dist_land (sd) | 1 | 120 | 21 | 89 | 1.01 | 2.68 | 5.01 | 1.94 |

**Table S1.5:**
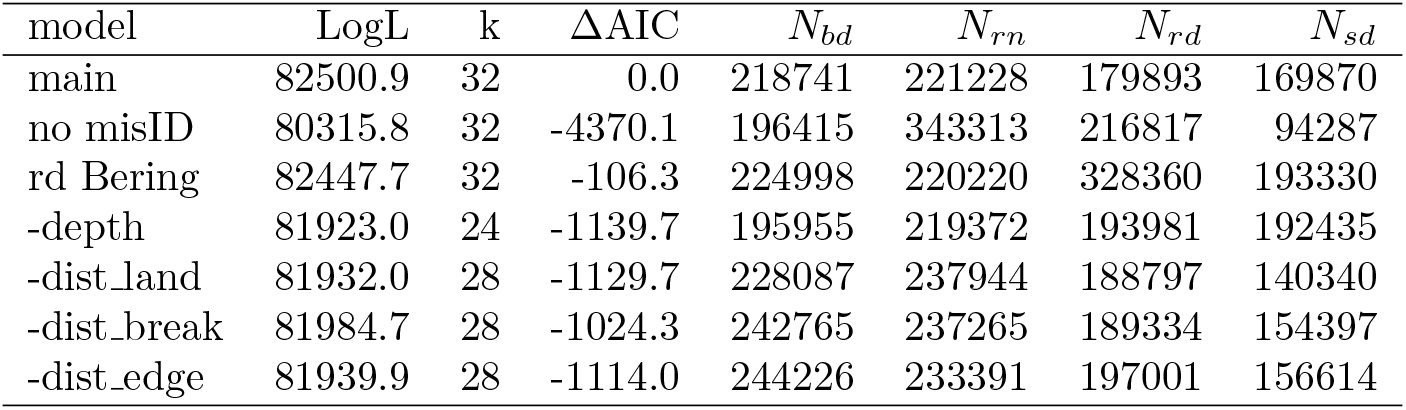
Results of sensitivity runs for 2013 surveys in the Sea of Okhotsk. The first column gives sensitivity run name, where “main” is the preferred model presented in the main paper, “no misID” is a model without species misidentification, and “rd Bering” uses alternative predictions of ringed seal availability. For the remaining models, ‘-’ and ‘+’ indicate whether a particular covariate is added or subtracted from the main model, and (sp) gives the species for which a covariate was added or substracted (bd: bearded; rn: ribbon; rd: ringed; sd: spotted). For each sensitivity run, we present the optimized log likelihood (LogL), the number of regression parameters (*k*), the difference in AIC score for each model compared to the “main” model, and abundance estimates for each species (Nbd - Nsd). The AIC score only included the number of regression parameters in the parameter count; positive values indicate increased predictive performance relative to our preferred model.

| model | LogL | k | ΔAIC | $N_{bd}$ | $N_{rn}$ | $N_{rd}$ | $N_{sd}$ |
| --- | --- | --- | --- | --- | --- | --- | --- |
| main | 82500.9 | 32 | 0.0 | 218741 | 221228 | 179893 | 169870 |
| no misID | 80315.8 | 32 | -4370.1 | 196415 | 343313 | 216817 | 94287 |
| rd Bering | 82447.7 | 32 | -106.3 | 224998 | 220220 | 328360 | 193330 |
| -depth | 81923.0 | 24 | -1139.7 | 195955 | 219372 | 193981 | 192435 |
| -dist_land | 81932.0 | 28 | -1129.7 | 228087 | 237944 | 188797 | 140340 |
| -dist_break | 81984.7 | 28 | -1024.3 | 242765 | 237265 | 189334 | 154397 |
| -dist_edge | 81939.9 | 28 | -1114.0 | 244226 | 233391 | 197001 | 156614 |

**Table S1.6:** Extrapolation metrics associated with sensitivity runs for 2013 surveys in the Sea of Okhotsk. The first column gives sensitivity run name, where “main” is the preferred model presented in the main paper, “no misID” is a model without species misidentification, and “rd Bering” uses alternative predictions of ringed seal availability. For the remaining models, ‘-’ and ‘+’ indicate whether a particular covariate is added or subtracted from the main model, and (sp) gives the species for which a covariate was added or substracted (bd: bearded; rn: ribbon; rd: ringed; sd: spotted). For each sensitivity run, we present the number of spatial cells for which abundance of a given species was predicted to be greater than the maximum predicted for a sampled cell *n_species_*, as well as the ratio of the maximum prediction in unsampled cells to the maximum prediction in sampled cells (Λ*_species_*).

| model | $n_{bd}$ | $n_{rn}$ | $n_{rd}$ | $n_{sd}$ | $\Lambda_{bd}$ | $\Lambda_{rn}$ | $\Lambda_{rd}$ | $\Lambda_{sd}$ |
| --- | --- | --- | --- | --- | --- | --- | --- | --- |
| main | 0 | 0 | 1 | 3 | 1.00 | 1.00 | 1.00 | 1.05 |
| no misID | 0 | 2 | 0 | 1 | 1.00 | 1.06 | 1.00 | 1.09 |
| rd Bering | 0 | 0 | 1 | 1 | 1.00 | 1.00 | 1.04 | 1.07 |
| -depth | 0 | 1 | 0 | 11 | 1.00 | 1.05 | 1.00 | 2.10 |
| -dist_land | 0 | 1 | 0 | 0 | 1.00 | 1.04 | 1.00 | 1.00 |
| -dist_break | 2 | 0 | 0 | 0 | 1.05 | 1.00 | 1.00 | 1.00 |
| -dist_edge | 0 | 2 | 0 | 0 | 1.00 | 1.07 | 1.00 | 1.00 |

## Supplementary Information 2

### Supplementary figures

**Figure S2.1:**
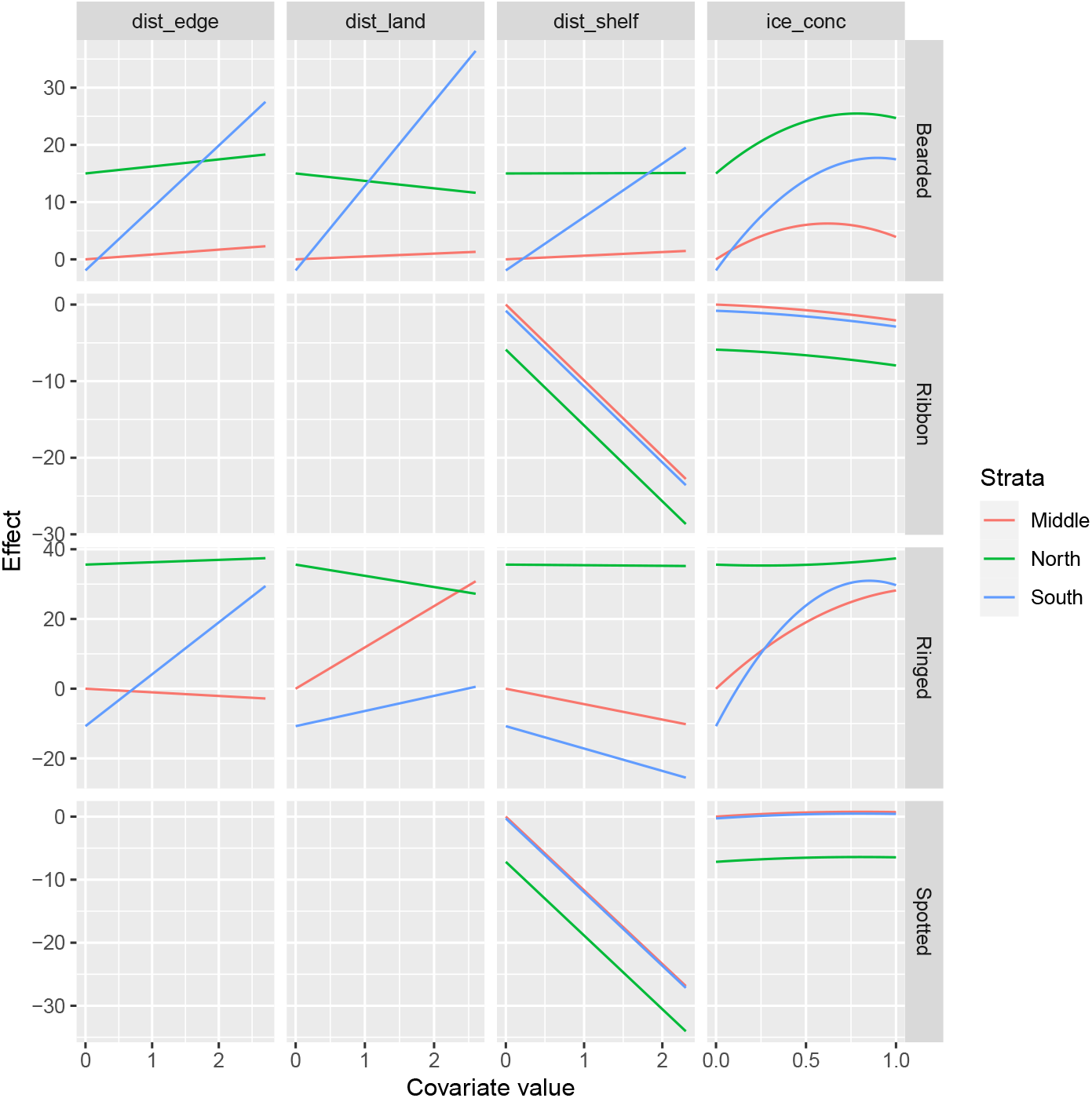
Effects of environmental and physiographic covariates on seal abundance in the western Bering Sea in 2012 on the scale of the linear predictor (i.e., logit scale). “Ice” represents the proportion of a grid cell with sea ice, and the remaining values represent distance to certain features (which were calculated in projected space and standardized to their mean, thus lacking meaningful units). In particular, “dist edge” represents distance to ice edge, “dist land” represents distance to land, and “dist shelf” represents distance to the closest 1000 m isobath. Covariates are measured at the centroid of each grid cell.

**Figure S2.2:**
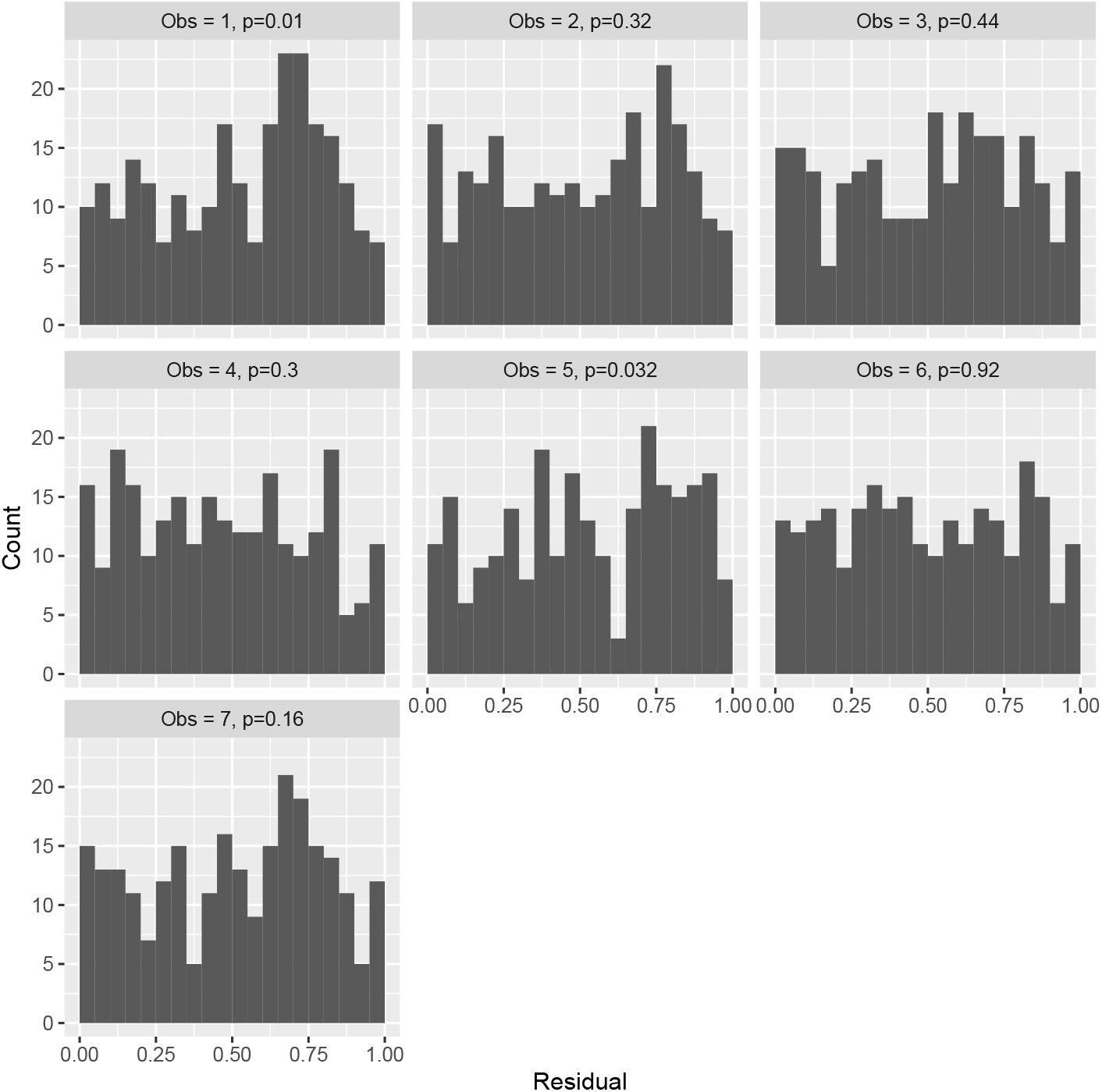
Randomized quantile residual diagnostics for our preferred “base” model fitted to count data obtained in 2012 aerial surveys in the western Bering Sea. Each histogram corresponds to counts of spotted seals (“Obs = 1”), ribbon seals (“Obs = 2”), bearded seals (“Obs = 3”), ringed seals (“Obs = 4”), unknown (but photographed) seal species (“Obs = 5”), pups (“Obs = 6”), and unphotographed seals (“Obs = 7”). In a well fitting model, each histogram would have a uniform distribution, an assumption which is tested with a *χ*^2^ discrepancy statistic (p-values in headers). In particular, we tend to under-predict medium-high counts of spotted seals.

**Figure S2.3:**
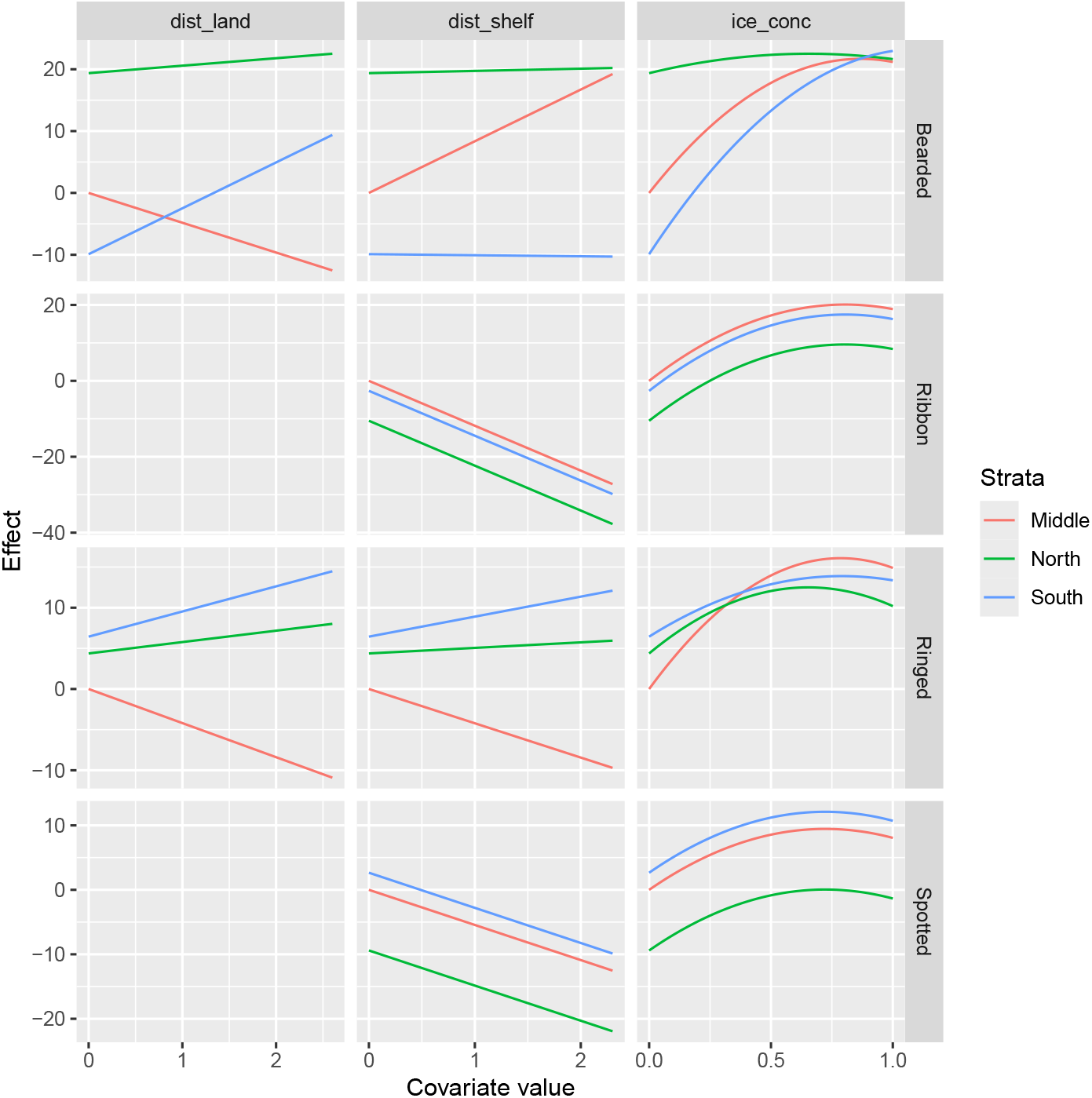
Effects of environmental and physiographic covariates on seal abundance in the western Bering Sea in 2013 on the scale of the linear predictor (i.e., logit scale). “Ice” represents the proportion of a grid cell with sea ice, and the remaining values represent distance to certain features (which were calculated in projected space and standardized to their mean, thus lacking meaningful units). In particular, “dist land” represents distance to land, and “dist shelf” represents distance to the closest 1000 m isobath. Covariates are measured at the centroid of each grid cell.

**Figure S2.4:**
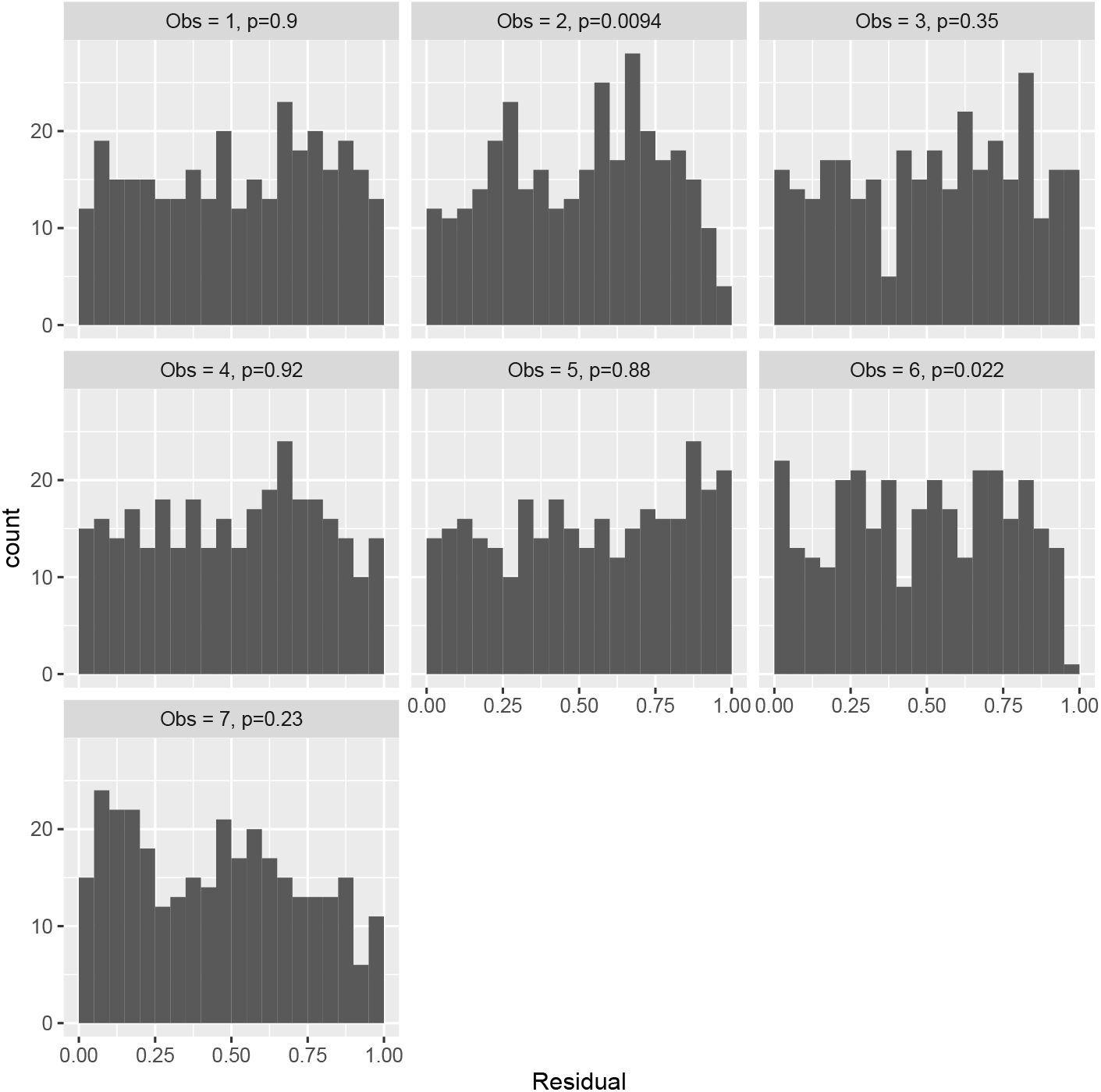
Randomized quantile residual diagnostics for our preferred “base” model fitted to count data obtained in 2013 aerial surveys in the western Bering Sea. Each histogram corresponds to counts of spotted seals (“Obs = 1”), ribbon seals (“Obs = 2”), bearded seals (“Obs = 3”), ringed seals (“Obs = 4”), unknown (but photographed) seal species (“Obs = 5”), pups (“Obs = 6”), and unphotographed seals (“Obs = 7”). In a well fitting model, each histogram would have a uniform distribution, an assumption which is tested with a *χ*^2^ discrepancy statistic (p-values in headers). In particular, we tend to over-predict medium-high counts of ribbon seals and under-predict very large counts of ribbon seals.

**Figure S2.5:**
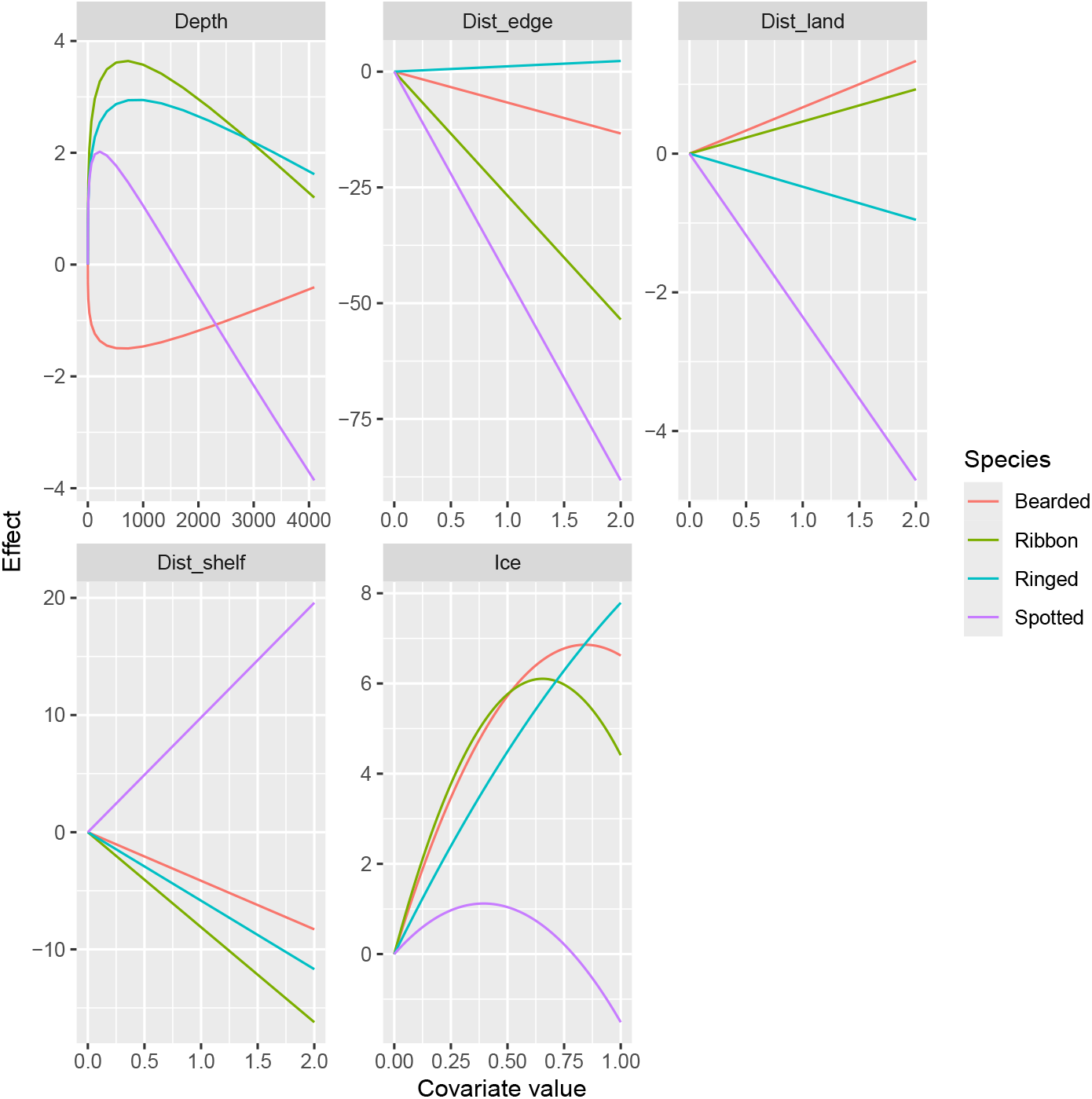
Effects of environmental and physiographic covariates on seal abundance in the Sea of Okhotsk on the scale of the linear predictor (i.e., logit scale). Depth is in meters, “Ice” represents the proportion of a grid cell with sea ice, and the remaining values represent distance to certain features (which were calculated in projected space and standardized to their mean, thus lacking meaningful units). In particular, “Dist_edge_” represents distance to ice edge, “Dist_land_” represents distance to land, and “Dist_shelf_” represents distance to the closest 200 m isobath. Covariates are measured at the centroid of each grid cell.

**Figure S2.6:**
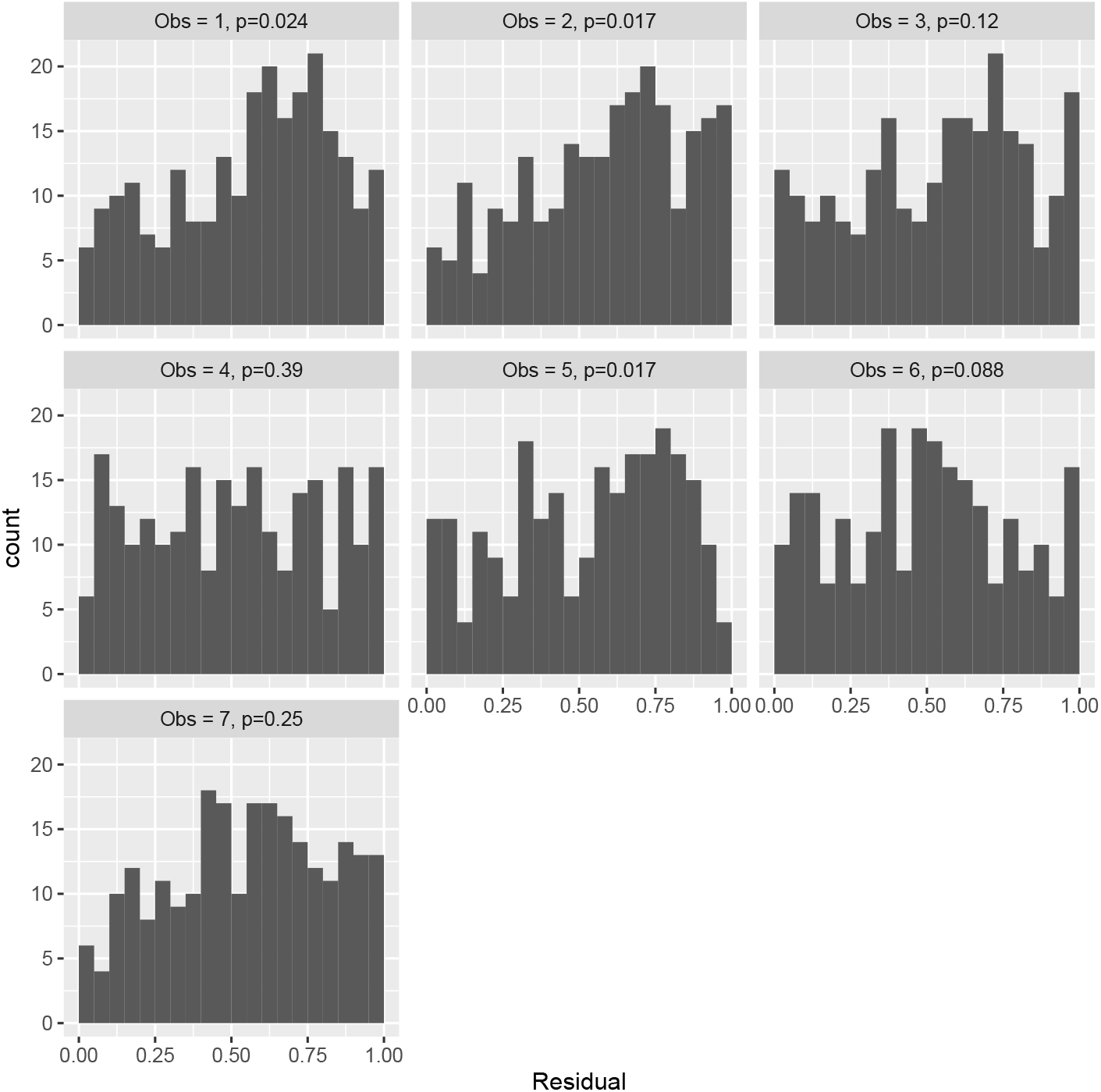
Randomized quantile residual diagnostics for our preferred “base” model fitted to count data obtained in 2013 aerial surveys in the Sea of Okhotsk. Each histogram corresponds to counts of spotted seals (“Obs = 1”), ribbon seals (“Obs = 2”), bearded seals (“Obs = 3”), ringed seals (“Obs = 4”), unknown (but photographed) seal species (“Obs = 5”), pups (“Obs = 6”), and unphotographed seals (“Obs = 7”). In a well fitting model, each histogram would have a uniform distribution, an assumption which is tested with a *χ*^2^ discrepancy statistic (p-values in headers). In particular, we tend to under-predict medium-high counts of spotted seals, high values of ribbon seals, and medium to high values of unknown (but photographed seal species).

## Notes

https://github.com/pconn/BOSSrussia

### Competing Interest Statement

The authors have declared no competing interest.

